# Mitochondrial Enzymes of the Urea Cycle Cluster at the Inner Mitochondrial Membrane

**DOI:** 10.1101/2020.05.18.102988

**Authors:** Nantaporn Haskins, Shivaprasad Bhuvanendran, Anna Gams, Tomas Kanholm, Kristen M. Kocher, Jonathan LoTempio, Kylie I. Krohmaly, Danielle Sohai, Nathan Stearrett, Erin Bonner, Mendel Tuchman, Hiroki Morizono, Jyoti K. Jaiswal, Ljubica Caldovic

## Abstract

Mitochondrial enzymes involved in energy transformation are organized into multiprotein complexes that channel the reaction intermediates for efficient ATP production. Three of the mammalian urea cycle enzymes: N-acetylglutamate synthase (NAGS), carbamylphosphate synthetase 1 (CPS1), and ornithine transcarbamylase (OTC) reside in the mitochondria. Urea cycle is required to convert ammonia into urea and protect the brain from ammonia toxicity. Urea cycle intermediates are tightly channeled in and out of mitochondria, indicating that efficient activity of these enzymes relies upon their coordinated interaction with each other perhaps in a multiprotein complex. This view is supported by mutations in surface residues of the urea cycle proteins that impair urea genesis in the patients but do not affect protein stability or catalytic activity. Further, we find one third of the NAGS, CPS1 and OTC proteins in liver mitochondria can associate with the inner mitochondrial membrane (IMM), and co-immunoprecipitate. Our *in silico* analysis of vertebrate NAGS proteins, the least abundant of the urea cycle enzymes, identified a region we call ‘variable segment’ present only in the mammalian NAGS protein. We experimentally confirmed that NAGS variable segment mediates the interaction of NAGS with CPS1. Use of Gated-Stimulation Emission Depletion (gSTED) super resolution microscopy showed that in situ, NAGS, CPS1 and OTC are organized into clusters. These results are consistent with mitochondrial urea cycle proteins forming a cluster instead of functioning either independently or in a rigid multienzyme complex.

## Introduction

Mitochondria are ATP producing organelles where multi-enzyme complexes catalyze reactions of the TCA cycle, fatty acid beta-oxidation and oxidative phosphorylation. To perform these diverse functions, mitochondria maintain the inner and outer mitochondrial membrane-bound compartments. These compartments are further spatially sub-organized into complexes to efficiently channel unstable and highly reactive intermediates between enzymes that catalyze consecutive reactions of the TCA cycle, oxidative phosphorylation and fatty acid beta-oxidation (Schmitt and An 2017). In addition to the compartments found ubiquitously in all mitochondria, the mammalian hepatocyte mitochondria contain three enzymes of the urea cycle, which converts ammonia into urea (Brusilow and Horwich 2001). Ammonia is a nitrogen waste product of protein catabolism and an extremely potent neurotoxin that can cause brain damage when its concentration in body fluids exceeds 50 μM (Brusilow and Horwich 2001). Therefore, the primary function of the urea cycle in mammals is to protect the central nervous system from the toxic effects of ammonia (Brusilow and Horwich 2001; Caldovic and Tuchman 2003). Six enzymes, N-acetylglutamate synthase (NAGS, EC 2.3.1.1), carbamylphosphate synthetase 1 (CPS1, EC 6.3.4.16), ornithine transcarbamylase (OTC, EC 2.1.3.3), argininosuccinate synthase (ASS EC 6.3.4.5), argininosuccinate lyase (ASL, EC 4.3.2.1) and arginase 1 (ARG1, EC 3.5.3.1) are required for the conversion of ammonia into urea (Brusilow and Horwich 2001). Because NAGS, CPS1 and OTC are located in the mitochondria and ASS, ASL and ARG 1 are cytoplasmic enzymes, two transporters, an ornithine/citrulline transporter (ORNT) and an aspartate/glutamate transporter, also known as citrin or ARALAR2, are also required for normal function of the urea cycle (Bradford and McGivan 1980; Kobayashi, et al. 1999; Brusilow and Horwich 2001). NAGS, CPS1 and OTC are also expressed in the intestinal mucosa where they catalyze formation of citrulline, which is a precursor for nitric oxide and arginine biosynthesis in mammals (Brusilow and Horwich 2001).

NAGS catalyzes formation of N-acetylglutamate (NAG), which is an essential allosteric activator of CPS1 in mammals. The amino acid sequence of the mammalian NAGS consists of three regions with different degrees of conservation: the mitochondrial targeting signal (MTS), the variable segment (VS), and the conserved segment (Caldovic, et al. 2002a). When mouse NAGS was expressed in cultured insect cells, the pre-protein was imported into the mitochondria and processed at two sites. Removal of the MTS resulted in a mature NAGS (NAGS-M) while removal of the MTS and the variable segment resulted in conserved NAGS (NAGS-C) (Morizono, et al. 2004). Recombinant NAGS-M and NAGS-C both catalyze the formation of NAG and are activated by arginine (Caldovic, et al. 2006).

CPS1 catalyzes the formation of carbamyl phosphate from ammonia, ATP, and bicarbonate, while OTC catalyzes the production of citrulline from carbamyl phosphate and ornithine (Brusilow and Horwich 2001). Citrulline is exported into cytoplasm via ORNT and converted into ornithine and urea by ASS, ASL and ARG1. Ornithine re-enters the urea cycle upon import into mitochondria by ORNT and urea is excreted by the kidneys (Brusilow and Horwich 2001). CPS1 is presumed to be the rate-limiting enzyme of ureagenesis because increased protein catabolism due to either high protein diet or breakdown of cellular proteins does not result in the accumulation of downstream urea cycle intermediates (Waterlow 1999).

NAGS, CPS1 and OTC have been considered to be soluble matrix proteins (Clarke 1976; Raijman 1976; Raijman and Jones 1976; Shigesada and Tatibana 1978; Bendayan and Shore 1982; Sonoda and Tatibana 1983; Hamano, et al. 1988). However, existing evidence suggests that these enzymes are not entirely freely distributed in the mitochondrial matrix but instead interact with each other and the inner mitochondrial membrane (IMM). Subcellular fractionation of rat liver mitochondria has revealed that CPS1 and OTC interacts with the IMM (Powers-Lee, et al. 1987), which is complemented by electron microscopy data showing that OTC is also closely associated with the IMM (Yokota and Mori 1986). Studies using isotopic tracers and isolated mitochondria have shown channeling of urea cycle intermediates from CPS1 to OTC (Cohen, et al. 1992) and from ASS to ASL to arginase 1 (Cheung, et al. 1989). Moreover, patients with urea cycle defects who receive a liver transplant continue to need supplementation with arginine (Tuchman 1989). This outcome of liver transplant suggests that due to tight channeling of intermediates between urea cycle enzymes arginine, an intermediate of urea cycle and a protein building block, does not leave transplanted liver, necessitating its continued supplementation.

The above properties of urea cycle suggest that mitochondrial urea cycle enzymes cluster allowing compartmentalization of urea cycle in the mitochondria. However, structural details of the urea cycle enzymes that allow such interaction and its clinical relevance remain unknown. Using a combination of protein structural analysis, mapping of patient mutations, liver mitochondrial fractionation, co-immunoprecipitation of urea cycle enzymes, and super-resolution microscopy we provide structural evidence that NAGS, CPS1 and OTC enzymes interact and form clusters in the mitochondria, providing direct evidence in support of urea cycle being yet another mitochondrial function that relies upon compartmentalization by the formation of protein super-complex.

## Materials and Methods

### Ethics Statement

Experimental procedures involving animals were approved by the Institutional Animal Care and Use Committee of the Children’s National Medical Center.

### Determination of the solvent accessible surface area and conservation

Crystal structures 5dot, 5dou and 1oth were used to calculate relative solvent accessible surface area (SASA) for the apo and liganded CPS1, and OTC trimer structures after removal of heteroatoms and water molecules. SASA of each amino acid was calculated with the Shrake and Rupley dot method (Shrake and Rupley 1973) as described by Ho (http://boscoh.com/protein/calculating-the-solvent-accessible-surface-area-asa.html) and using mesh density 9600. A custom Python script (https://github.com/morizono/pdbremix) was used to calculate SASA for each residue. The same method was used to calculate maximal SASA for amino acid using polypeptide in which each of the 20 amino acids is flanked by a glycine residue (Supplementary File 1); this polypeptide was modeled as b-strand using VEGA 3.1.1 (Pedretti, et al. 2002). Relative SASA was calculated by dividing SASA of each amino acid with its maximal SASA.

Conservation of amino acids was determined from the alignment of either 233 homologs of human CPS1 (Supplementary File 2) or 270 homologs of human OTC (Supplementary File 3) from vertebrates and multicellular invertebrates. Protein sequences were collected from the NCBI non-redundant protein sequence database using Protein BLAST (Altschul, et al. 1990; Altschul, et al. 1997), default parameters (word size 6, expected threshold 10, scoring matrix BLOSSUM62, gap existence 11, and gap extension 1) and sequences of human CPS1 and OTC as queries. Clustal Omega (Madeira, et al. 2019) was used for multiple protein sequence alignment. Conservation of surface residues that are mutated in patients with CPS1 and OTC deficiencies was determined as percent of either 233 CPS1 or 270 OTC homologs that have the same amino acid as human protein at that position.

### Identification and Computational analysis of VS sequences

Protein sequences of vertebrate NAGS were collected using Blastp to query vertebrate proteins in either NCBI nr or UniProt databases with human and zebrafish NAGS (Caldovic, et al. 2002a; Caldovic, et al. 2014). Default parameters (word size 6, gap opening and extension penalties 11 and 1, respectively, and BLOSUM62 scoring matrix) were used for the search, which resulted in 90 mammalian NAGS sequences (Supplementary file 4) and 61 NAGS sequences from fish, amphibians and reptiles (Supplementary File 5). The most likely translation initiation site for each NAGS sequence was determined by inspection of protein alignments with the corresponding genomic sequences and with human and zebrafish NAGS, performed using ClustaW in MEGA7 (Kumar, et al. 2018); amino acids encoded by predicted exons located upstream of the exon that corresponds to exon 1 in human and zebrafish *NAGS* genes were removed. The boundaries of the VS were defined as sequences between the mitochondrial protein peptidase (MPP) cleavage site and the beginning of sequence homology with vertebrate-like N-acetylglutamate synthase-kinase from *Xanthomonas campestris* (XcNAGS-K), which does not have MTS and VS (Qu, et al. 2007). Sequence alignments with mouse NAGS, which has experimentally determined MPP cleavage site (Caldovic, et al. 2010), as well as MitoPorotII (Claros and Vincens 1996) and MitoFates (Fukasawa, et al. 2015) were used for prediction of MPP cleavage sites in NAGS sequences. The C-termini of VS were determined by sequence alignments of NAGS sequences with XcNAGS-K using ClustalW in MEGA7. The lengths, proline content and sequence identities of VS were determined using MEGA7. WebLOGO3 (Crooks, et al. 2004) was used to visualize VS sequence alignments that were generated with ClustalW in MEGA7.

### Fractionation of Rat Liver Mitochondria

Fractionation of mitochondria was carried out as described previously (Powers-Lee, et al. 1987). Briefly, mitochondria were purified from donated rat livers using differential centrifugation (Graham 2001). Purified mitochondria were resuspended in 5mM Tris HCl, 250mM Sucrose, 1mM EDTA, pH 7.2, and subjected either to three rounds of freezing and thawing, or treatment with 0.12 mg of digitonin per mg of mitochondrial protein to remove the outer mitochondrial membrane. Mitoplasts were separated from outer membrane vesicles using centrifugation at 9000xg for 10 min. The mitoplasts were resuspended in 20mM Hepes Buffer, pH 8.0 and sonicated to obtain inner mitochondrial membrane vesicles. The vesicles were treated with increasing concentrations of Triton X-100 (0, 0.1, 0.5 and 1.0%) for 30 min. at room temperature, followed by pelleting of the membranes by ultracentrifugation at 144,000xg for 60 min, washing three times with 20mM Hepes, pH 8.0, and re-suspension in the same buffer. The amount of NAGS in each mitochondrial fraction was determined using immunoblotting with the primary antibody raised against recombinant mouse NAGS at 1:5,000 dilution and HPRT-conjugated donkey anti-rabbit secondary antibody (Pierce) at 1:50,000 dilution. NAGS bands were visualized using SuperSignal West Pico kit (Pierce) according to the manufacturer’s instructions. The amounts of CPS1 and OTC in each mitochondrial fraction were determined using immunoblotting with primary antibodies raised against CPS1 or OTC at 1:5,000 dilution, followed by the HPRT-conjugated secondary antibody at 1:10,000 dilution. CPS1 and OTC were visualized using ECL Western Blotting Substrate (Pierce) according to the manufacturer’s instructions. The intensity of each band was measured using a GS-800 Calibrated Densitometer (Bio-Rad) and the Quantity One software package (Bio-Rad). Mitochondrial fractions were probed with antibodies raised against mitochondrial markers of the IMM, mitochondrial matrix and outer mitochondrial membrane: CoxIV (Abcam) at 1:5,000 dilution, Grp75 (Stressgen) at 1:1,000 dilution and VDAC (Pierce) at 1:1,000 dilution (Da Cruz, et al. 2003), (Rardin, et al. 2008; Rardin, et al. 2009). Filters were then probed with the HPRT-conjugated secondary antibody (Bio-Rad). The CoxIV, Grp75, and VDAC bands were visualized using ECL Western Blotting Substrate (Pierce).

### Cloning of Recombinant Mouse Variable Segments

Mouse variable segment (mVS) coding sequence was subcloned using pNS1 plasmid (Caldovic, et al. 2002b) as a template and primers 5’-GGG ACA TAT GCT CAG CAC CGC CAG GGC TCA C-3’ and 5’-AGG TGG ATC CTT ATT ATT ACC AGT GGC GTG CTT CC-3’ which amplify the sequence between codons 49 and 117 of the mouse NAGS preprotein coding sequence. The amplification conditions were: initial denaturation at 95°C for 3 min., followed by 25 cycles of denaturation at 95°C for 30 sec., annealing at 60°C for 30 sec. and extension at 72°C for 30 sec., and final extension at 72°C for 5 min. using *Pfu* Turbo Hotstart DNA polymerase (Stratagene). This amplification product was cloned into pCR4Blunt-TOPO (Invitrogen) producing TOPOmVS. The correct coding sequence was confirmed by DNA sequencing. Plasmid TOPOmVS was cleaved with *Nde*I and *Bam*HI sites and subcloned into pET15b to create pET15bmVS.

The amino acid sequence of the reversed variable segment (revVS) was generated by reversing the amino acid sequence of mVS. The amino acid sequence of shuffled variable segment (shVS) was generated by dividing the sequence of mVS in the middle, then inter-digitating the amino acid sequences of the two halves. The coding sequences of revVS and shVS, including three stop codons at their 3’ ends and *Nde*I and *BamH*I restriction sites at the 5’- and 3’-ends, were chemically synthesized as mini-genes and inserted into pIDTSMART-KAN plasmid (Integrated DNA Technologies) followed by subcloning into pET15b bacterial expression vector to create pET15brevVS and pET15bshVS plasmids.

### Recombinant Protein Purification

Recombinant NAGS was purified as described previously (Caldovic, et al. 2006; Haskins, et al. 2008). Briefly, plasmid pET15bmNAGS-M (Caldovic, et al. 2006) was used for overexpression of mouse NAGS-M in E. coli. Pelleted cells were resuspended in Buffer A (50mM potassium phosphate, 500mM KCl, 20% glycerol, 10mM b-mercaptoethanol, 0.006%Triton X-100, 1% acetone, pH 7.5) containing 10 mM imidazole and lysed with 40 mM n-octyl-b-d-glucopyranoside. Cell lysate was loaded onto Ni-NTA agarose colum and recombinant NAGS-M was eluted with Buffer A containing 250 mM imidazole.

Recombinant mVS, revVS and shVS were purified from cultures of transformed *Escherichia coli* C41(DE3) cells that were induced with the Overnight Express Autoinduction Kit System 1 (Novagen). Cells were pelleted and resuspended in Buffer A containing 10mM imidazole. Lysozyme and phenylmethylsulfonyl fluoride were added to the final concentrations of 1mg/ml and 0.1mM respectively. The cells were lysed with 40mM n-octyl-b-D-glucopyranoside. DNAseI and RNAseA (0.1-0.5mg/ml lysate) in 5mM MgCl_2_ were added to remove nucleic acids by incubation at room temperature for 30 min. Cell lysate was cleared by centrifugation at 25,000xg for 30 min at 4°C. A nickel-affinity column (GE Healthcare) was equilibrated with buffer A containing 10mM imidazole. Cleared lysate was loaded onto the column at a flow rate of 0.3ml/min. The column was washed with Buffer A containing 50, 125, 250 and 500mM imidazole. The variable segments eluted between 250 and 500mM imidazole. The protein size and purity were verified by Commassie blue staining following SDS-PAGE on the 16.5% Tris-Tricine Gel (Bio-Rad).

### Mass Spectrometry Peptide Sequencing of Mouse Variable Segments

To confirm the identity of the purified mouse variable segments, they were excised from the 16.5% Tris-Tricine Gel and subjected to rapid, in-gel trypsin digestion (Shevchenko, et al. 2006). The fragments were analyzed using mass spectrometry on an Applied Biosystems Voyager 4700 MALDI TOF/TOF mass spectrometer.

### Co-immunoprecipitation

Mouse liver mitochondria were purified from donated tissue using differential centrifugation (Graham 2001) and lysed with PBS containing 2% CHAPS (Stankiewicz, et al. 2005). Mitochondrial lysate was diluted to 1 mg/ml protein for immunoprecipitation with antibodies against OTC and CPS1 and 5 mg/ml protein for immunoprecipitation with anti-NAGS antibodies. Mitochondrial lysates were mixed with magnetic beads (Invitrogen) cross-linked to primary antibodies against NAGS, CPS1 or OTC according to the manufacturer’s instructions. Following incubation at 20°C for 10 min, the beads were washed five times with PBS containing 0.05% Triton X-100. Protein complexes were eluted with ImmunoPure IgG Elution Buffer (Pierce). Protein concentration in each elution fraction was measured using protein assay dye reagent concentrate (Bio-Rad) according to the manufacturer’s instructions. Between 0.5 and 1 *μ*g of immunoprecipitated proteins were resolved using SDS-PAGE, and probed with primary antibodies raised against NAGS, CPS1 or OTC followed by the HPRT-conjugated secondary antibody. NAGS was visualized using SuperSignal West Pico kit (Pierce), and CPS1 and OTC were visualized using ECL Western Blotting Substrate (Pierce).

In experiments measuring competition between NAGS-M and the recombinant mVS, mitochondrial lysate was diluted to a protein concentration of 2 mg/ml, mixed with the mVS, revVS or shVS in a 1:1 (v/v) ratio, and added to magnetic beads (Invitrogen) cross-linked to primary antibodies against CPS1. Depending on the experiment, the molar excess of recombinant variable segment peptides relative to CPS1 was between 10- and 30-fold, based on estimates of the reported abundance of CPS1 in the liver mitochondria (Raijman 1976; Cohen, et al. 1982; Sonoda and Tatibana 1983; Wang, et al. 2019a). Immunoprecipitation was carried out as described above. The intensities of NAGS-M bands were measured using a GS-800 Calibrated Densitometer and Quantity One software (Bio-Rad).

### Confocal and gSTED microscopy

Confocal and Gated Stimulated Emission Depletion (GSTED) microscopy were performed as described previously (Bhuvanendran, et al. 2014; Salka, et al. 2017). Imaging was performed using the Leica TCS SP8 microscope equipped with a white light laser, two depletion lasers, acousto-optical beam splitter (AOBS) and hybrid detectors. Single labeling of all the confocal and GSTED samples was done using Alexa Fluor 647 while the double-labeled samples also had Alexa Fluor 532.

An HC PL APO CS2 100x/1.40 Oil objective was used to acquire confocal images. Alexa Fluor 532 was excited using 515 nm laser line and the emission was collected on a hybrid detector with the AOBS set to 520–590 nm whereas the Alexa Fluor 647 was excited using 645 nm laser line and the emission between 650–720 nm was collected.

12-bit GSTED images with pixel size less than 30 nm were acquired using the HC PL APO CS2 100x/1.40 Oil objective. Samples with Alexa Fluor 647 fluorophores were excited at 645 nm and depleted with 775 nm laser. The emission was collected between 650 nm to 720 nm with a time gating of 0.3 ns to 6.0 ns. In the double-labeled samples, sequential stack for Alexa Fluor 532 was acquired using a 515 nm excitation and 660 nm depletion. The time-gated emission between 2.2 ns and 6.0 ns was collected with the AOBS set from 520 nm to 590 nm.

These confocal and STED images were deconvolved with Huygens Professional version 17.04 (Scientific Volume Imaging, http://svi.nl). Further image analysis, including intensity plots, were done using MetaMorph Premier version 7.7.0 (Molecular Devices, www.metamorph.com).

## Results and Discussion

### Comparative analysis of CPS1 and OTC surface residues whose mutations disrupt ureagenesis and cause disease

More than half of CPS1 and OTC deficiency cases are caused by missense mutations (Haberle, et al. 2011; Caldovic, et al. 2015). Due to the tight channeling of CPS1 and OTC intermediates in the mitochondria (Cheung, et al. 1989; Cohen, et al. 1992) we reasoned that some of the disease-causing missense variants of CPS1 and OTC may disrupt their interaction. To assess this, we calculated relative solvent accessible surface area (SASA) of CPS1 and OTC amino acids and mapped the positions of amino acids whose replacements cause CPS1 and OTC deficiencies.

Human CPS1 pre-protein is 1500 amino acids long. 1352 and 1422 amino acids are visible in the apo and liganded state, respectively (de Cima, et al. 2015). Of these, 556 and 525 amino acids have more than 25% of their surface area exposed to solvent in the apo and liganded states, respectively, which we refer to as surface residues/amino acids (Levy 2010). Of the 161 missense mutations that cause CPS1 deficiency (Table S1), 22 mutations affect residues that are on the surface of both apo and liganded CPS1, 18 affect surface residues of the apo CPS1, and three affect surface residues of the liganded CPS1 (Table S2). Biochemical properties of the p.Y389C, p.A438T, p.T471N, p.R721Q, p.K875E, p.E1255D, p.R1262Q, p.R1262P, p.C1327R, p.R1371L, p.P1439L, p.T1443A and p.Y1491H recombinant CPS1 revealed that mutations of these surface residues result in destabilization, decreased enzymatic activity, and/or decreased affinity for NAG (Pekkala, et al. 2009; Diez-Fernandez, et al. 2013; Diez-Fernandez, et al. 2014; Diez-Fernandez, et al. 2015) and Table S2). Inspection of the CPS1 3D-structure (de Cima, et al. 2015) strongly suggested that p.R174W, p.D358H, p.K450E, p.G628D, p.R780H, p.R814W, p.Q1059R, and p.R1089C replacements could disrupt electrostatic interactions with nearby charged residues (Table S2). The p.R1317W and p.G1333E replacements affect amino acids in the T’-loop and may disrupt substrate channeling between CPS1 active sites (de Cima, et al. 2015), while replacement of the A438 with proline could affect flexibility of and hydrogen bonding within the T-loop (de Cima, et al. 2015; Gao, et al. 2015). Disruption of the proteinprotein interactions with OTC and/or NAGS could explain CPS1 deficiency caused by the remaining 14 missense mutations that affect CPS1 surface residues (Figure 1 and Table S2). Additionally, the p.M1148, p.A1155E and p.A1155V affect residues that are not visible in the CPS1 crystal structures suggesting that this region of CPS1 might be disordered. Disordered protein domains have been shown to mediate protein-protein interactions (Hsu, et al. 2012) and amino acid replacements within such domains could disrupt binding of the interacting partners. Functional importance of the M1148, A1155 and the 14 surface amino acids whose replacements cause CPS1 deficiency was evaluated by determining their conservation in CPS1 homologs from 233 animal species (Supplementary File 2). Seven of these residues are 100% conserved, and additional six are over 85% conserved in CPS1 homologs, while residues that correspond to less conserved A640, H1045, and R1228 have similar size and/or chemical properties in most species (Table 1).

**Figure 1.**
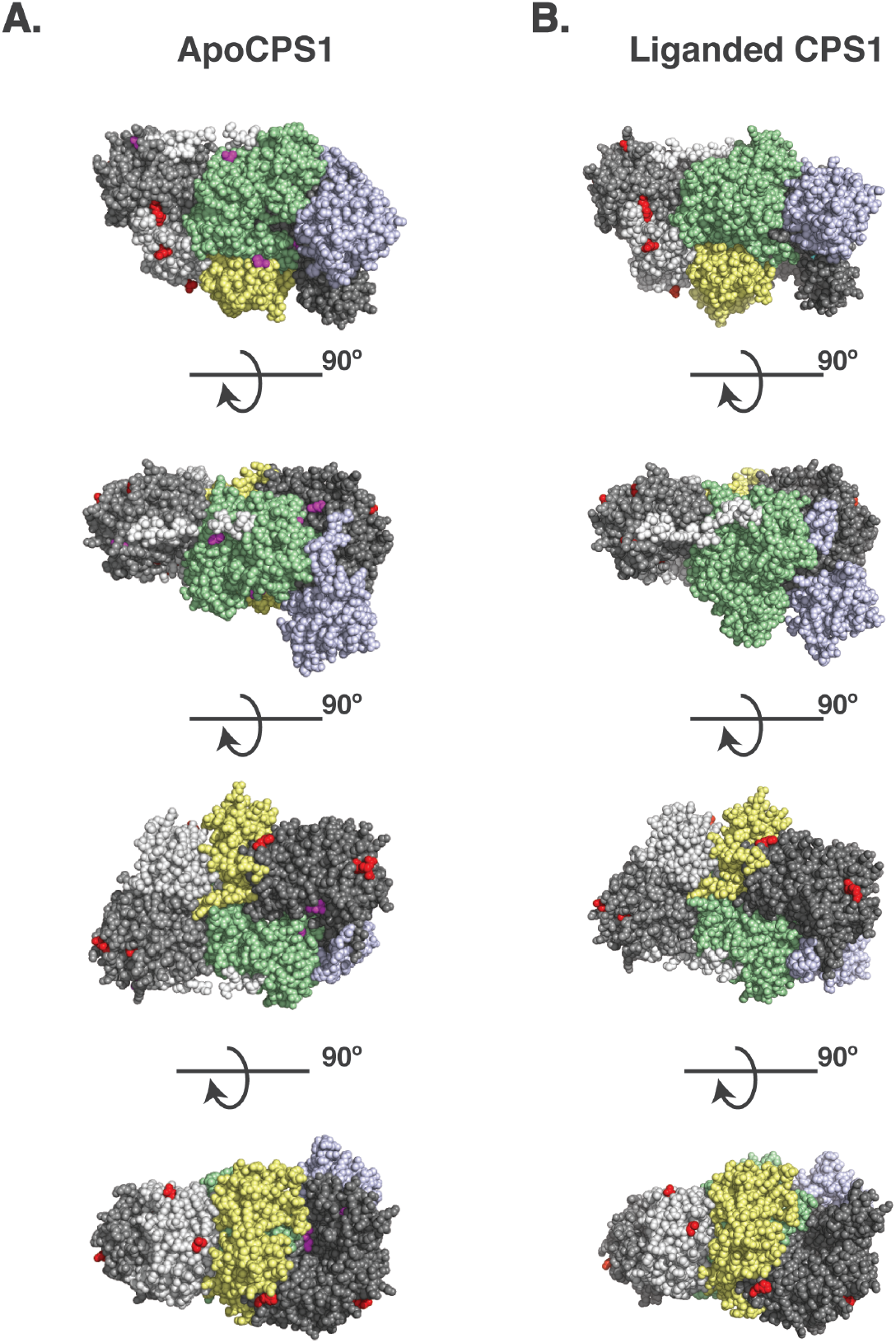
Surface residues in the apo (A) and liganded CPS1 (B) whose replacements cause CPS1 deficiency. Disease causing mutations in the *CPS1* gene were mapped to 3-dimensional structures of the apo CPS1 (5dot) and liganded CPS1 (5dou) enzymes. **A**. Eight surface residues in apo CPS1 whose replacements cause CPS1 deficiency are shown in red. Five surface residues that are exposed in apo CPS1 only are shown in magenta. **B**. Eight surface residues in liganded CPS1 whose replacements cause CPS1 deficiency are shown in red. One surface residue that is exposed only in liganded CPS1 is shown in cyan.

**Table 1.** Conservation of CPS1 surface residues whose replacements cause CPS1 deficiency.

| Residue ID | Conservation Score<sup>a</sup> | Amino Acids in CPS1 and CPS3 from other Species |  |  |  |  |  |
| --- | --- | --- | --- | --- | --- | --- | --- |
|  |  | Mammals | Birds | Reptiles | Amphibians | Fish | Invertebrates |
| A43 | 89 | A | A/V/M | A | A | A/T/G | A/T/S/P |
| R233 | 100 | R | R | R | R | R | R |
| K280 | 97 | K | K | K | K | K/N/Q | K/Q/E/T |
| A304 | 98 | A/T | A | A | A | A/V/S | A |
| G401 | 87 | G/E/K/A/N/P | G/E/K | G/E | G | G/E/Q/W | G/K |
| A640 | 69 | A/S | A | A | A | A/I/V | A/S/C |
| P846 | 97 | P/A | P | P | P | P | P/A |
| I986 | 100 | I | I | I | I | I | I |
| G987 | 100 | G | G | G | G | G | G |
| H1045 | 42 | H/Q | Q | Q | Q | H/Q/R/E | H/Q/E/S |
| V1141 | 100 | V | V | V | V | V | V |
| *M1148<sup>c</sup>* | *92* | *M* | *M* | *M* | *L* | *M* | *L* |
| *A1155<sup>d</sup>* | *100* | *A* | *A* | *A* | *A* | *A* | *A* |
| R1228 | 69 (90<sup>b</sup>) | R/W/Q | R/K/E | R/K | K | R/W/H/Q/C | R/K/Q/A |
| I1254 | 100 | I | I | I | I | I | I |
| D1274 | 100 | D | D | D | D | D | D |

<sup>a</sup>Conservation was determined based on the alignment of CPS1 and CPS3 protein sequences from 233 animals.
<sup>b</sup>Conservation of positively charged amino acids, R or K, in this position.
<sup>c</sup>Residue has 12% SASA in the apoCPS1 structure and is not visible in the liganded structure.
<sup>d</sup>Residue not visible in the apoCPS1 and liganded CPS1 structures.

Mature human OTC protein is 322 amino acids long and the functional enzyme is a trimer (Shi, et al. 1998; Caldovic, et al. 2015). In each subunit 125 residues have over 25% of their surface area accessible to solvent. Of the 265 missense mutations that cause OTC deficiency (Table S3), 51 are replacements of 35 surface amino acids. Deleterious effects of the p.R40H, p.T49P, p.A102P, p.H255P, p.Q270P, p.L349P, p.G269E, p.G269R, p.K221N, p.K289D and p.K289N replacements can be explained by their effects on either *OTC* mRNA splicing or protein folding, stability, and catalytic properties of the mutant protein (Table S4). Functional importance of the remaining 25 surface residues (Figure 2) was evaluated by determining their conservation in OTC proteins from 270 animal species (Supplementary File 3). 15 surface amino acids associated with deleterious missense mutations are over 85% conserved in OTC homologs and the remaining 10 are replaced in most species by amino acids with similar size and/or chemical properties (Table 2).

**Figure 2.**
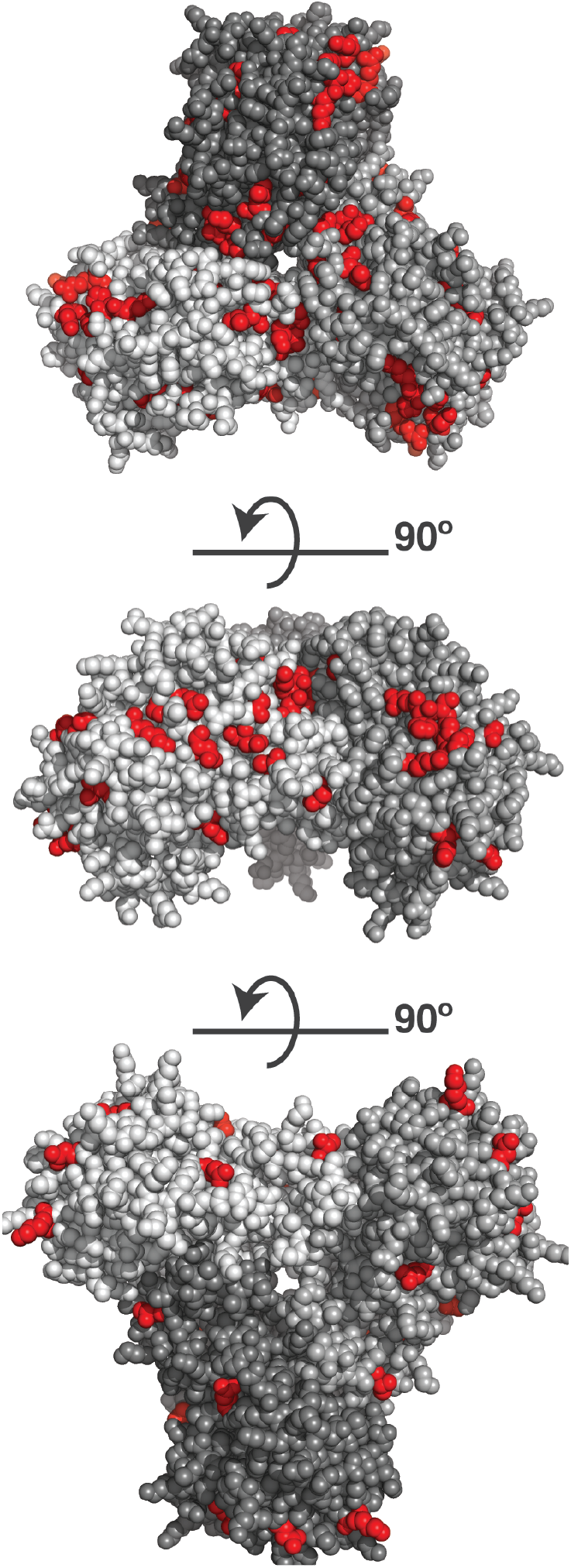
Surface residues in the OTC and whose replacements cause OTC deficiency. Disease causing mutations in the *OTC* gene were mapped to 3-dimensional structures of the human OTC (1oth); 25 surface residues whose replacements cause OTC deficiency are shown in red.

**Table 2.**
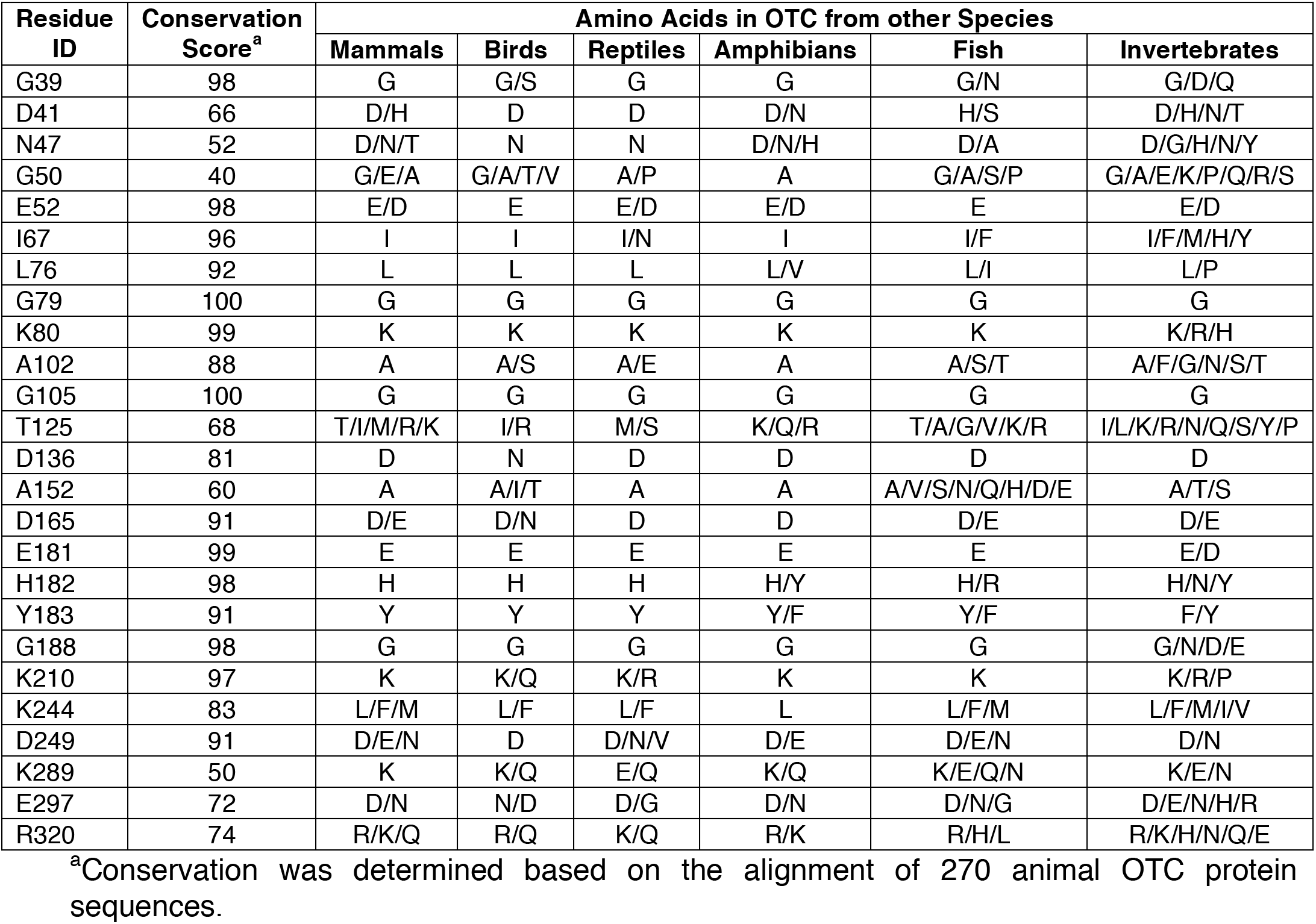
Conservation of OTC surface residues whose replacements cause OTC deficiency.

Conservation of surface amino acids in CPS1 and OTC that are mutated in patients with urea cycle deficiency suggests potential importance of these residues for the functioning of the urea cycle. Given that these amino acids are located away from the catalytic and substrate binding sites, and do not interact with other amino acids within the proteins, we hypothesize that these surface amino acids may mediate protein-protein interactions between CPS1 and OTC.

### Protein-protein Interactions Between NAGS, CPS1 and OTC in the Liver Mitochondria

To test the above hypothesis, we used co-immunoprecipitation to probe the interaction between NAGS, CPS1 and OTC. Each protein was immunoprecipitated using the protein-specific antibody, while non-specific antibody was used as a negative control. Purified recombinant NAGS (Figure 3A) and total liver proteins (Figures 3B and 3C) were used as positive controls. Use of NAGS-specific antibody allowed co-immunoprecipitation of both CPS1 and OTC (Figures 3B and 3C). Concordantly, CPS1-specific antibody co-immunoprecipitated NAGS and OTC (Figure 3A and 3C), and OTC-specific antibody co-immunoprecipitated NAGS and CPS1 (Figure 3A and 3B). The ability of each of the proteins to co-immunoprecipitate the other two urea cycle proteins expressed at endogenous levels in the liver offers direct evidence in support of the ability of each of these proteins to interact with the other urea cycle proteins in the liver mitochondria.

**Figure 3.**
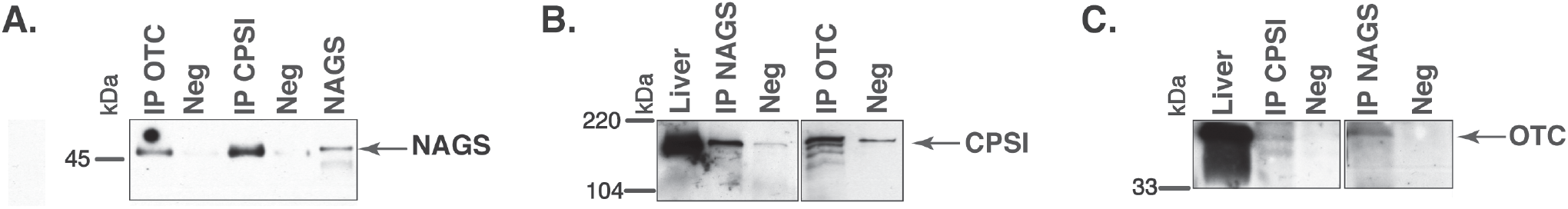
Co-immunoprecipitation of NAGS, CPSI and OTC from the mouse liver. Anti-NAGS (IP NAGS lane), anti-CPSI (IP CPSI lane) and anti-OTC (IP OTC lane) antibodies were used for immunoprecipitation. Non-specific IgG (Neg) was used as a negative control. The precipitated proteins were probed with antibodies against NAGS (A), CPSI (B) and OTC (C). Either 0.5 ng of recombinant NAGS (panel A) or 5 *μ*g (panels B and C) of liver proteins (Liver lane) were used as positive controls. Between 0.5 and 1 *μ*g of immunoprecipitated proteins were resolved in IP NAGS, IP CPSI and IP OTC lanes.

In view of the ability of the urea cycle proteins to interact we next examined the protein region responsible for this promoting these interactions and focused on NAGS. We hypothesized that interactions between NAGS, CPS1 and OTC are crucial for increasing the efficiency of the urea cycle. While NAGS is present in all organisms, its efficient activity is crucial for land dwelling organisms such as mammals as their survival requires highly efficient disposal of nitrogenous waste (Cohen 1963; Mommsen and Walsh 1989; Haskins, et al. 2008). Inversion of allosteric effect of L-arginine from inhibition to activation of NAGS in land dwelling tetrapods is a feature of NAGS that enabled efficient ureagenesis (Haskins, et al. 2008). Thus, we compared mammalian NAGS proteins with NAGS from fish, amphibians and reptiles; birds lack *NAGS* genes (Haskins, et al. 2008). Variable segments (VS) from mammalian NAGS proteins are more conserved, longer and have higher proline content than VS from fish, amphibian and reptile NAGS proteins. (Figure 4 and Table 3). Proline-rich protein segments can form extended, poly-proline type II helices that mediate protein-protein interactions (Rubin, et al. 2000; Kelly, et al. 2001; Ball, et al. 2005). In view of the co-IP of NAGS with CPS1 and presence of the proline-rich interaction domain only in NAGS from mammals, we examined if the VS mediates interaction of mammalian NAGS (Caldovic, et al. 2002a) with CPS1 and whether this interaction may be important for NAGS function in the mammals.

**Figure 4.**
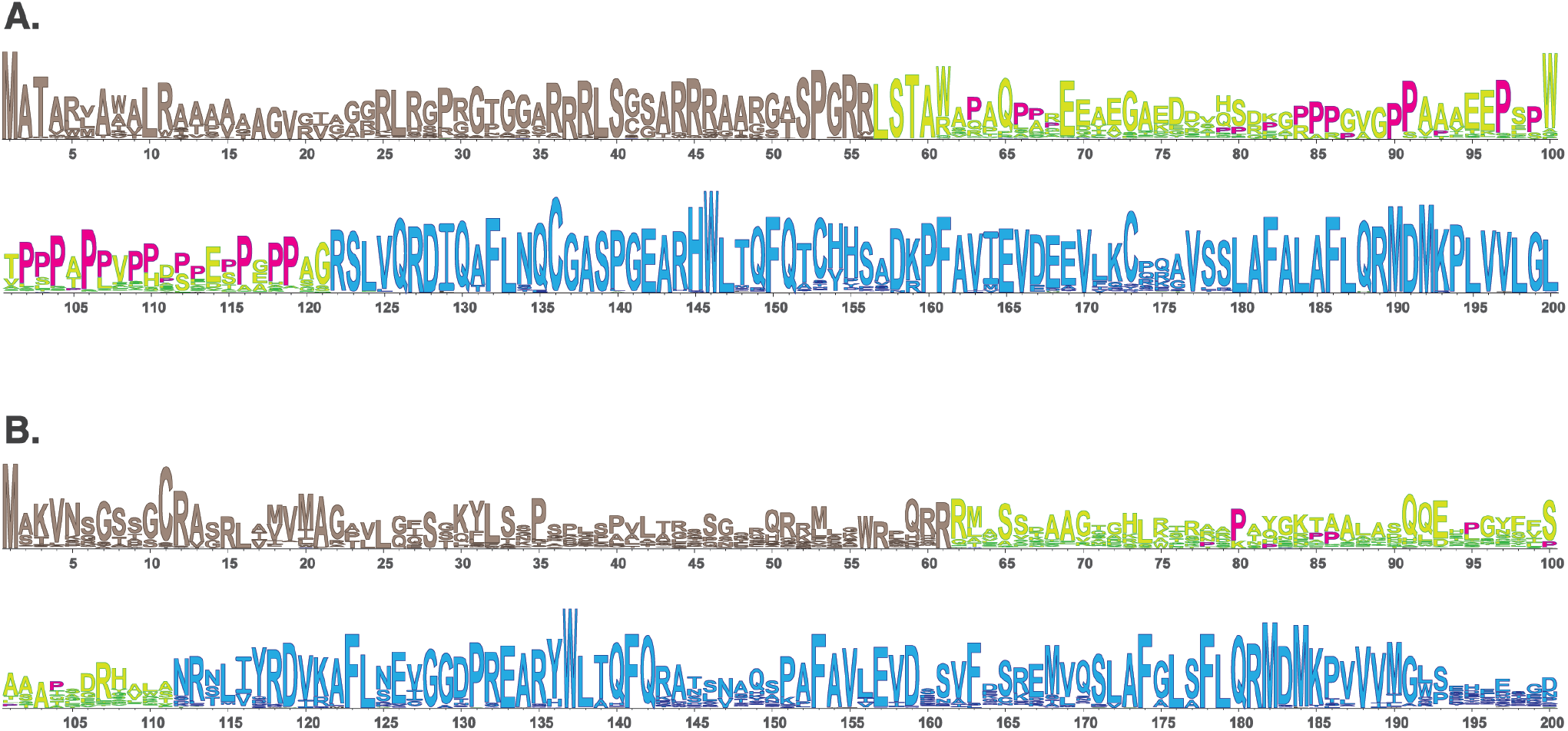
LOGO alignments of the N-terminal portions of NAGS from 90 mammals **(A)** and 61 fish, amphibians and reptiles **(B)**. NAGS is present in most vertebrates but its efficient activity is crucial for survival of land dwelling organisms such as mammals that require highly efficient urea cycle. To examine protein sequence features associated with NAGS specialization in aquatic and land dwelling vertebrates we compared alignments of NAGS from mammals and fish, amphibians and reptiles. There are tree regions with differing degrees of conservation: N-terminal mitochondrial targeting signal, followed by VS, and conserved domain. Mammalian VS has higher proline content and greater sequence conservation than the VS from fish, amphibians and reptiles. Prolines are shown in magenta. **Tan** – mitochondrial targeting sequence. **Yellow/lime green** – variable segment (VS). **Blue** – conserved domain.

**Table 3.** Properties of vertebrate VS.

|  | **Mammals** | **Fish, Amphibians and Reptiles** |
| --- | --- | --- |
| **VS Length** | 37-64 | 31-40 |
| **VS Proline Content** | 17.4 – 32.3% | 2.3 – 11.1 % |
| **VS Sequence Identity** | 24 – 84% | 34 – 60% |

To assess whether NAGS variable segment mediates NAGS-CPS1 interactions, we used purified recombinant mouse variable segment (mVS) to determine if it competes with mitochondrial NAGS for co-immunoprecipitation with CPS1. Two peptides with identical amino acid composition as mVS but altered in their amino acid sequence: reverse mouse variable segment (revVS) and a shuffled mouse variable segment (shVS) polypeptide were used as negative controls (Figure 5A). Recombinant mVS, revVS and shVS, tagged with poly-histidine at the N-terminus, were overexpressed in *E. coli* and purified using nickel affinity chromatography. Purified mVS, revVS and shVS migrated as 9.5 kDa bands, which are in close agreement with their predicted molecular weights of 9675 Da (Figure 5B). Mass spectrometry peptide fingerprinting and peptide sequencing of purified mVS, revVS and shVS were used to confirm their sequences (Supplementary Figure S1). Recombinant mVS, revVS or shVS were added to liver mitochondrial lysate followed by co-immunoprecipitation using the anti-CPS1 antibody and probing with the anti-NAGS antibody. Total liver protein was used as positive control for immunoblotting. Separate co-immunoprecipitations were performed using non-specific IgG antibodies, revVS (IP CPS1+revVS) or shVS (IP CPS1+shVS). Figure 5C illustrates the results. Overall, the addition of recombinant mVS reduced the amount of mNAGS that co-immunoprecipitated with the CPS1 by at least 5-fold. Neither revVS nor shVS displaced mNAGS from CPS1.

**Figure 5.**
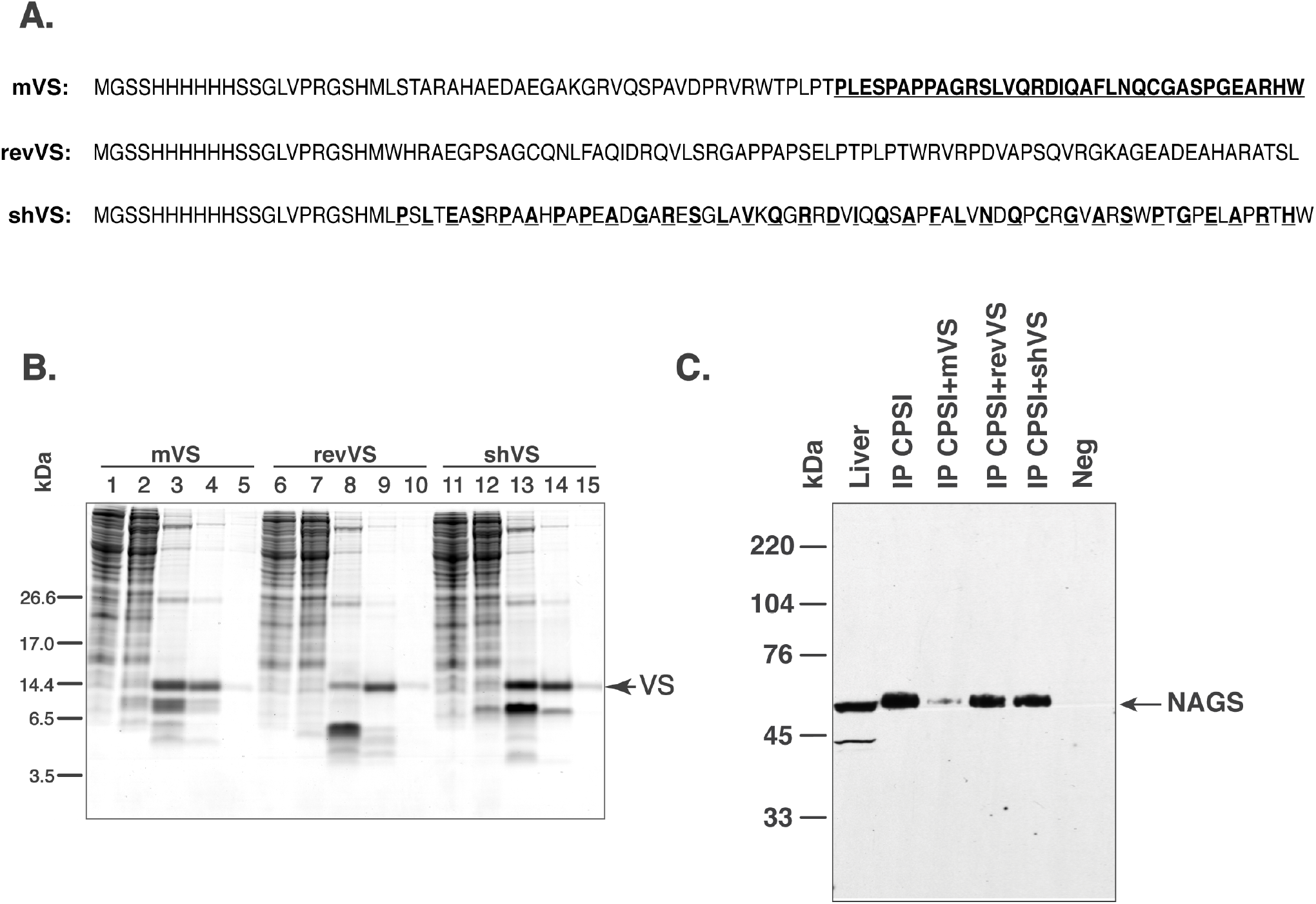
Competition between NAGS and mouse variable segment (mVS) for co-IP with CPSI. Recombinant mVS was overexpressed and purified from *E. coli*. **A**. Amino acid sequences of recombinant mouse variable segment (mVS), reversed (revVS) and shuffled (shVS) variable segments. Amino acids from the C-terminal half of mVS (bold and underlined typeface) were interdigitated with amino acids from the N-terminal half to create shVS. **B**. Purification of mVS, rVS and shVS. Lanes 1, 6 and 11 – flow trough. Lanes 2, 7 and 12 – wash with 50 mM imidazole. Lanes 3, 8 and 13 – elution with 125 mM imidazole. Lanes 4, 9, and 14 – elution with 250 mM imidazole. Lanes 5, 10 and 15 – elution with 500 mM imidazole. **C**. Anti-CPSI antibodies were used for immunoprecipitation of mitochondrial proteins. The precipitated proteins were probed with anti-mNAGS antibodies. Precipitation with non-specific antibodies (Neg), revVS (IP CPSI+revVS) and shVS (IP CPSI+shVS) were negative controls. Total liver proteins (Liver) were positive control.

Our immunoprecipitation experiments suggest stable interaction between NAGS, CPS1 and OTC. The CPS1 monomer and OTC trimer, which are the active forms of these two enzymes, are present in approximately a 10:1 molar ratio in the liver mitochondria (Raijman 1976; Cohen, et al. 1982; Wang, et al. 2019a), while NAGS is approximately one thousand times less abundant than CPS1 (Sonoda and Tatibana 1983; Wang, et al. 2019a). The stoichiometry of the NAGS-CPS1-OTC complex may reflect the abundance of each protein in the liver mitochondria. It is possible that the NAGS is tethered to one molecule of CPS1 via VS and provides NAG to a number of neighboring CPS1 molecules. Another possibility is a dynamic interaction between NAGS, CPS1 and OTC may result in more than one complex of the three proteins.

### Distribution of NAGS, CPS1 and OTC in Liver Mitochondria

While above studies identify interaction between the urea cycle enzymes, they do not identify the location in the mitochondria where these interactions occur. Earlier fractionation studies found that CPS1 and OTC exist both in the mitochondrial matrix with a significant fraction loosely attached to IMM (Powers-Lee, et al. 1987). Thus, we used CPS1 and OTC as positive controls to examine NAGS distribution in the soluble and membrane-associated fractions of rat liver mitochondria. Glucose related protein 75 (Grp75), subunit IV of the cytochrome c oxidase (Cox IV), and voltage–dependent anion channel (VDAC) were used as additional reporters to mark the soluble matrix, inner and outer mitochondrial membranes, respectively (Da Cruz, et al. 2003; Rardin, et al. 2008; Rardin, et al. 2009). These proteins confirmed the purity of the mitochondrial fractions (lanes 2 and 6 in Figure 6). Approximately 25% of NAGS partitioned with the membrane fraction (lane 2 in Figure 6), while 75% of NAGS was in soluble fraction (lane 6 in Figure 6).

**Figure 6.**
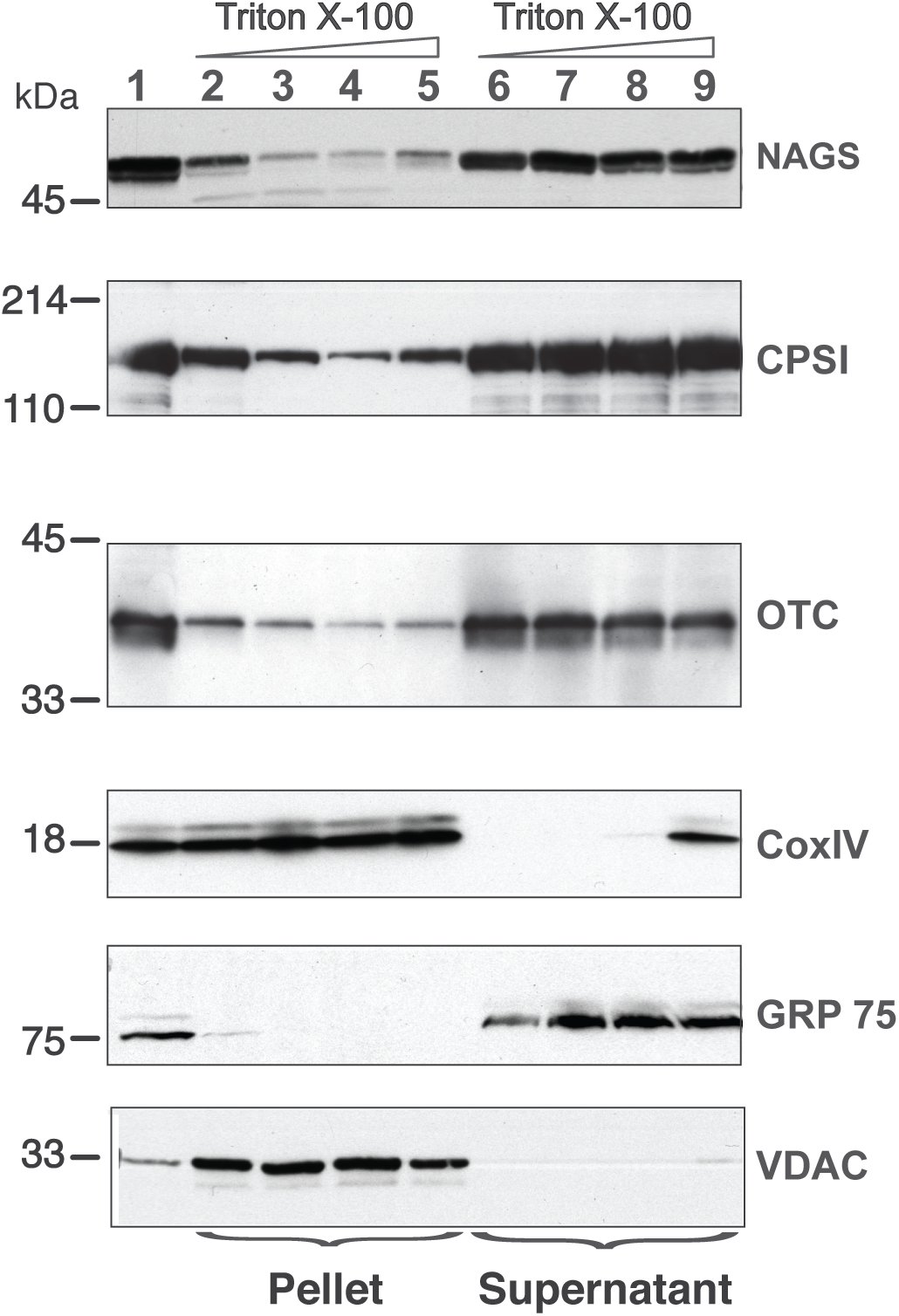
Distribution of NAGS, CPSI and OTC in the liver mitochondria. Increasing amounts of TritonX-100 were added to sonicated mitoplasts; proteins associated with the membrane (pellet) were separated from the soluble proteins (supernatant) and probed with the anti-NAGS, anti-CPSI, anti-OTC, anti-CoxIV, anti-Grp75 and anti-VDAC. 30 *μ*g of liver mitochondrial proteins were used as positive control for NAGS and 2 *μ*g of mitochondrial proteins was used as positive control for OTC, CPSI, COXIV, Grp75 and VDAC. Lane 1 – liver mitochondrial proteins, lanes 2 and 6 – 0% TritonX-100, lanes 3 and 7 – 0.1% TritonX-100, lanes 4 and 8 – 0.5%, TitonX-100, lanes 5 and 9 – 1% TritonX-100.

The addition of 0.1% Triton X-100 reduced the membrane-associated fraction of NAGS to about 10% (lane 3 in Figure 6). Increasing detergent concentrations, which dissociates more tightly bound proteins from membranes, did not increase solubilization of NAGS. Addition of Triton X-100 did not result in solubilization of Cox IV and VDAC (lanes 3-5 in Figure 6), while Grp75 remained soluble under the same experimental conditions (lanes 7-9 in Figure 6). Similar to our results with NAGS, and consistent with previous studies (Powers-Lee, et al. 1987), 30% of CPS1 and 20% of OTC were associated with the IMM (Figure 6). Again, similar to NAGS, even at the highest detergent concentrations, CPS1 and OTC enzymes did not completely dissociate from the membranes. This suggests that part of the NAGS, OTC and CPS1 protein complex in the mitochondrial membrane exists in a detergent (0.1% Triton X-100)-resistant compartment. To further confirm that membrane association of these enzymes is not an artefact of outer mitochondrial membrane (OMM) removal by freezing and thawing, we carried out mitochondrial fractionation after solubilization of OMM with digitonin. This independently confirmed the distribution of NAGS, CPS1 and OTC at the IMM and in the matrix (Supplementary Figure S2).

Above biochemical studies established that of NAGS, CPS1 and OTC partition between IMM and mitochondrial matrix, which perhaps regulates the reserve capacity for ureagenesis in the matrix. These results suggest that the urea cycle enzymes exist in the mitochondria in one of the two extreme cases outlined in Figure 7A. To directly visualize membrane association of the urea cycle enzymes in situ and assess if they are present in a cluster we used gated Stimulated Emission Depletion (gSTED) super-resolution microscopy to localize NAGS, CPS1 and OTC in primary mouse hepatocytes. If these urea cycle enzymes are uniformly distributed in the mitochondrial matrix, then we would expect lack of any co-localized protein clusters (Figure 7A, left), whereas interactions of the three proteins with each other at the IMM, we would result in observation of NAGS, CPS1, and OTC enzyme clusters (Figure 7A, right). Confocal microscopy established the mitochondrial localization of the urea cycle enzymes (Figure 7B), while use of gSTED microscopy identified that, in addition to some diffuse localization in the mitochondrial matrix, these enzymes are all detectable in clusters away from the mitochondrial matrix. These clusters were 100-150 nm in size, and thus small enough to be below the resolution limit of confocal microscopy (Figure 7C-G). By co-immunostaining we observed that these clusters were not formed by individual proteins, but contained miltiple urea cycle enzymes – NAGS and OTC (Figure 7H, I), and NAGS and CPS1 (Figure 7J, K). These *in situ* results support the interaction observed between NAGS, CPS1 and OTC by our biochemical co-localization studies. While some NAGS and OTC co-clustered as indicated by overlapping fluorescence peaks for green (NAGS) and red (OTC) pixels (Figure 7I), we did observe NAGS and OTC clusters that did not co-localize (Figure 7H, I). We observed that such independent clusters were greater for the abundant urea cycle enzyme, CPS1 (Figure 7J, K). Thus, *in situ* superresolution visualization approach identified that, while a proportion of the more abundant urea cycle enzymes (OTC, CPS1) co-localize with NAGS clusters, these proteins can also exist in independent clusters (red only peaks in Figures 7I, K), and can be detected more diffusely along the IMM or in the mitochondrial matrix (Figure 7H-K). NAGS, the less abundant and potentially regulatory urea cycle protein, was only detected in clusters that are limited to co-clusters with CPS1, as indicated by the spatial overlap of green (NAGS) and red (CPS1/OTC) peaks in Figures 7I and 7K.

**Figure 7.**
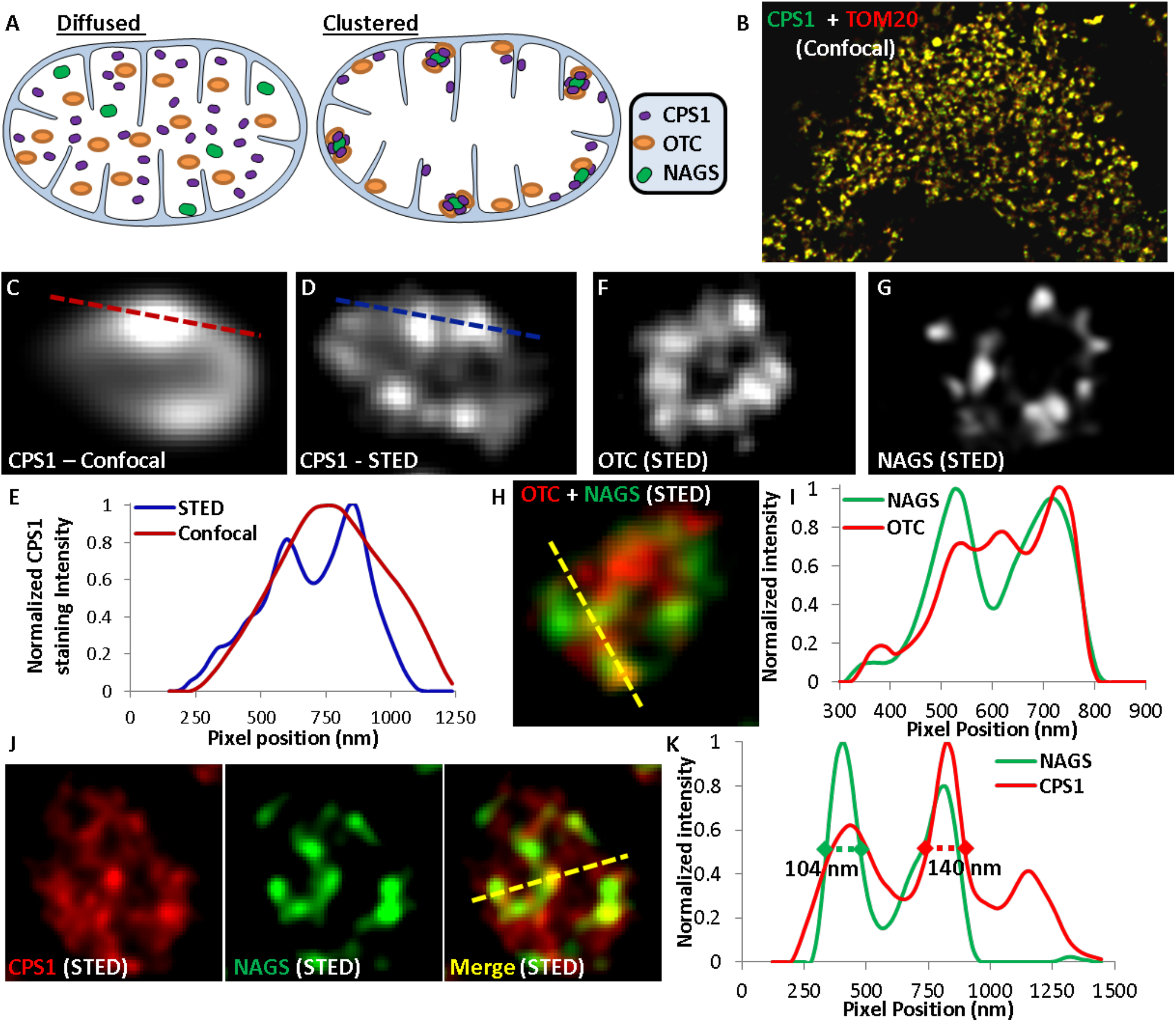
Use of super-resolution microscopy to determine in situ nanoscale organization of urea cycle enzymes. (**A**) Schematic representing the possible distribution of urea cycle enzymes in the mitochondria. (**B**) Confocal images showing localization of CPS1 and the mitochondrial protein TOM20 labeled with Alexa Fluor 532 and Alexa Fluor 647 respectively. (**C**) Confocal and (**D, F, G**) GSTED images showing individual mitochondria in mouse hepatocytes immuno-stained for (**C, D**) CPS1, (**F**) OTC, (**G**) NAGS, (**H**) OTC (Alexa Fluor 647) and NAGS (Alexa Fluor 532), (**J**) CPS1 (Alexa Fluor 647) and NAGS (Alexa Fluor 532). Single labeling in images **C, D, F, and G** was done with Alexa Fluor 647. (**E, I, K**) Normalized pixel intensity plot for the dotted lines marked in the corresponding confocal or STED images.

Our co-immunoprecipitation and super resolution microscopy data demonstrate that NAGS, CPS1 and OTC interact with each other and are present in a cluster along the mitochondrial membrane. While co-immunoprecipitation of the three proteins suggests stable interactions, the large difference in their abundance (Raijman 1976; Cohen, et al. 1982; Sonoda and Tatibana 1983; Wang, et al. 2019a) raises questions regarding stoichiometry of such complex. Our data, taken together with large differences in abundance of NAGS, CPS1 and OTC, are consistent with clustering of the three proteins at the IMM, similar to enzymes for biosynthesis of coenzyme Q and purines (An, et al. 2008; Chan, et al. 2015; French, et al. 2016; Subramanian, et al. 2019) Enzyme clustering provides metabolic advantages without the need for evolution of complimentary protein-protein interfaces (Sweetlove and Fernie 2018). Theoretical modeling and experiments with engineered proteins in *E. coli* show that clustering of enzymes that catalyze consecutive reactions of a metabolic pathway can increase flux through the pathway by 100-fold; this is achieved through increased local concentrations of enzymes and metabolites in the cluster (Castellana, et al. 2014). Another advantage of enzyme clusters is protection of unstable, highly reactive metabolites, such as carbamyl phosphate, from hydrolyzing or reacting with other metabolites or surface residues of proteins (Sweetlove and Fernie 2018). The identity of surface amino acids could be important for their ability to cluster. We identified replacements of highly conserved CPS1 and OTC surface residues that may disrupt their clustering and effective catalysis of citrulline formation, which would result in accumulation of ammonia. This is similar to disease-causing mutations at the surface of adenylosuccinate lyase that disrupt formation of purinosome, a cluster of enzymes in *de novo* purine synthesis (Baresova, et al. 2012). The mitochondrial space appears to be organized into sub-compartments containing enzymes of different metabolic pathways. Enzymes in the Krebs’ cycle, electron transport, and fatty acid oxidation pathways form multiprotein complexes (Sumegi and Srere 1984; Robinson and Srere 1985; Robinson, et al. 1987; Velot, et al. 1997; Wu and Minteer 2015; Bulutoglu, et al. 2016; Liang, et al. 2018; Wang, et al. 2019b; Xia, et al. 2019) while mitochondrial urea cycle enzymes appear to form clusters that can accommodate large differences in the abundance of NAGS, CPS1, and OTC. Biochemical characterization of the interactions between NAGS, CPS1 and OTC is needed to establish the driving force for their clustering. This driving force could be electrostatic, hydrophobic, or both and may require presence of the IMM phospholipids. Additional super resolution imaging will provide information about organization of the NAGS-CPS1-OTC cluster and determine its spatial relationship to enzymes in other mitochondrial metabolic pathways. This is important because production of each urea molecule consumes three ATP molecules and efficient ureagenesis may rely on the proximity of CPS1 and protein complexes of the ATP producing pathways. Understanding of the driving force for clustering of NAGS, CPS1 and OTC will enable development of new treatments designed to promote and stabilize their interactions when one of the enzymes is defective.

## Author Contributions

NH – Co-immunoprecipitation of NAGS, CPS1 and OTC, overexpression and purification of recombinant mVS, revVS and shVS, competition of mVS and NAGS for co-immunoprecipitation with CPS1, mitochondrial fractionation and immunoblotting, immunofluorescence of NAGS, CPS1 and OTC clusters.

SB – Immunofluorescence and image analysis of NAGS, CPS1 and OTC clusters.

AG, TK, KMK, JL, KK, DS, NS, and EB – Carried out bioinformatic analysis of the NAGS variable segment ass ad a class project in a graduate-level course taught by LC.

MT – Design of biochemical studies and critical revisions of the manuscript.

HM – Calculation of the relative SASA for CPS1 and OTC amino acids.

JKJ – Design of the immunofluorescence studies, data analysis and critical review of the manuscript.

LC – Mapping of the CPS1 and OTC patient mutations, design of biochemical studies, preparation and revision of the manuscript

## Acknowledgments

We would like to thank Dr. Kristy Brown and Dr. Yetrib Hathout for their help with mass spectrometry and proteomics analyses, as well as Dr. Norma Allewell and Dr. Nicholas AhMew for helpful discussions and critical reading of this manuscript.

## References

Altschul, SF, Gish, W, Miller, W, Myers, EW and Lipman, DJ 1990 Basic local alignment search tool. J Mol Biol 215: 403–410

Altschul, SF, Madden, TL, Schaffer, AA, et al. 1997 Gapped blast and psi-blast: A new generation of protein database search programs. Nucleic Acids Res 25: 3389–3402

An, S, Kumar, R, Sheets, ED and Benkovic, SJ 2008 Reversible compartmentalization of de novo purine biosynthetic complexes in living cells. Science 320: 103–106

Ball, LJ, Kuhne, R, Schneider-Mergener, J and Oschkinat, H 2005 Recognition of proline-rich motifs by protein-protein-interaction domains. Angew Chem Int Ed Engl 44: 2852–2869

Baresova, V, Skopova, V, Sikora, J, et al. 2012 Mutations of atic and adsl affect purinosome assembly in cultured skin fibroblasts from patients with aica-ribosiduria and adsl deficiency. Hum Mol Genet 21: 1534–1543

Bendayan, M and Shore, GC 1982 Immunocytochemical localization of mitochondrial proteins in the rat hepatocyte. J Histochem Cytochem 30: 139–147

Bhuvanendran, S, Salka, K, Rainey, K, et al. 2014 Superresolution imaging of human cytomegalovirus VMIA localization in sub-mitochondrial compartments. Viruses 6: 1612–1636

Bradford, NM and McGivan, JD 1980 Evidence for the existence of an ornithine/citrulline antiporter in rat liver mitochondria. FEBS Lett 113: 294–298

Brusilow, SW and Horwich, AL 2001 Urea cycle enzymes CR Scriver, AL Beaudet, WS Sly and D Valle The metabolic & molecular bases of inherited disease McGraw-Hill, 1909–1963

Bulutoglu, B, Garcia, KE, Wu, F, Minteer, SD and Banta, S 2016 Direct evidence for metabolon formation and substrate channeling in recombinant tca cycle enzymes. ACS Chem Biol 11: 2847–2853

Caldovic, L, Abdikarim, I, Narain, S, Tuchman, M and Morizono, H 2015 Genotypephenotype correlations in ornithine transcarbamylase deficiency: A mutation update. J Genet Genomics 42: 181–194

Caldovic, L, Ah Mew, N, Shi, D, Morizono, H, Yudkoff, M and Tuchman, M 2010 N-acetylglutamate synthase: Structure, function and defects. Mol Genet Metab 100 Suppl 1: S13–19

Caldovic, L, Haskins, N, Mumo, A, et al. 2014 Expression pattern and biochemical properties of zebrafish N-acetylglutamate synthase. PLoS One 9: e85597

Caldovic, L, Lopez, GY, Haskins, N, et al. 2006 Biochemical properties of recombinant human and mouse N-acetylglutamate synthase. Mol Genet Metab 87: 226–232

Caldovic, L, Morizono, H, Gracia Panglao, M, et al. 2002a Cloning and expression of the human n-acetylglutamate synthase gene. Biochem Biophys Res Commun 299: 581–586

Caldovic, L, Morizono, H, Yu, X, et al. 2002b Identification, cloning and expression of the mouse n-acetylglutamate synthase gene. Biochem J 364: 825–831

Caldovic, L and Tuchman, M 2003 N-acetylglutamate and its changing role through evolution. Biochem J 372: 279–290

Castellana, M, Wilson, MZ, Xu, Y, et al. 2014 Enzyme clustering accelerates processing of intermediates through metabolic channeling. Nature biotechnology 32: 1011–1018

Chan, CY, Zhao, H, Pugh, RJ, et al. 2015 Purinosome formation as a function of the cell cycle. Proc Natl Acad Sci U S A 112: 1368–1373

Cheung, CW, Cohen, NS and Raijman, L 1989 Channeling of urea cycle intermediates in situ in permeabilized hepatocytes. J Biol Chem 264: 4038–4044

Clarke, S 1976 A major polypeptide component of rat liver mitochondria: Carbamyl phosphate synthetase. J Biol Chem 251: 950–961

Claros, MG and Vincens, P 1996 Computational method to predict mitochondrially imported proteins and their targeting sequences. Eur J Biochem 241: 779–786

Cohen, NS, Cheung, CW, Kyan, FS, Jones, EE and Raijman, L 1982 Mitochondrial carbamyl phosphate and citrulline synthesis at high matrix acetylglutamate. J Biol Chem 257: 6898–6907.

Cohen, NS, Cheung, CW, Sijuwade, E and Raijman, L 1992 Kinetic properties of carbamoyl-phosphate synthase (ammonia) and ornithine carbamoyltransferase in permeabilized mitochondria. Biochem J 282 (Pt 1): 173–180

Cohen, SS 1963 On biochemical variability and innovation. Science 139: 1017–1026

Crooks, GE, Hon, G, Chandonia, JM and Brenner, S, E. 2004 Weblogo: A sequence logo generator. Genome Res. 14: 1188–1190

Da Cruz, S, Xenarios, I, Langridge, J, Vilbois, F, Parone, PA and Martinou, JC 2003 Proteomic analysis of the mouse liver mitochondrial inner membrane. J Biol Chem 278: 41566–41571

de Cima, S, Polo, LM, Diez-Fernandez, C, et al. 2015 Structure of human carbamoyl phosphate synthetase: Deciphering the on/off switch of human ureagenesis. Sci Rep 5: 16950

Diez-Fernandez, C, Gallego, J, Haberle, J, Cervera, J and Rubio, V 2015 The study of carbamoyl phosphate synthetase 1 deficiency sheds light on the mechanism for switching on/off the urea cycle. J Genet Genomics 42: 249–260

Diez-Fernandez, C, Hu, L, Cervera, J, Haberle, J and Rubio, V 2014 Understanding carbamoyl phosphate synthetase (cps1) deficiency by using the recombinantly purified human enzyme: Effects of cps1 mutations that concentrate in a central domain of unknown function. Mol Genet Metab 112: 123–132

Diez-Fernandez, C, Martinez, AI, Pekkala, S, et al. 2013 Molecular characterization of carbamoyl-phosphate synthetase (CPS1) deficiency using human recombinant cps1 as a key tool. Hum Mutat 34: 1149–1159

French, JB, Jones, SA, Deng, H, et al. 2016 Spatial colocalization and functional link of purinosomes with mitochondria. Science 351: 733–737

Fukasawa, Y, Tsuji, J, Fu, SC, Tomii, K, Horton, P and Imai, K 2015 Mitofates: Improved prediction of mitochondrial targeting sequences and their cleavage sites. Mol Cell Proteomics 14: 1113–1126

Gao, M, Zhou, H and Skolnick, J 2015 Insights into disease-associated mutations in the human proteome through protein structural analysis. Structure 23: 1362–1369

Graham, JM 2001 Isolation of mitochondria from tissues and cells by differential centrifugation. Curr Protoc Cell Biol Chapter 3: Unit 3 3

Haberle, J, Shchelochkov, OA, Wang, J, et al. 2011 Molecular defects in human carbamoy phosphate synthetase i: Mutational spectrum, diagnostic and protein structure considerations. Hum Mutat 32: 579–589

Hamano, Y, Kodama, H, Yanagisawa, M, Haraguchi, Y, Mori, M and Yokota, S 1988 Immunocytochemical localization of ornithine transcarbamylase in rat intestinal mucosa. Light and electron microscopic study. J Histochem Cytochem 36: 29–35

Haskins, N, Panglao, M, Qu, Q, et al. 2008 Inversion of allosteric effect of arginine on n-acetylglutamate synthase, a molecular marker for evolution of tetrapods. BMC biochemistry 9: 24

Hsu, WL, Oldfield, C, Meng, J, et al. 2012 Intrinsic protein disorder and protein-protein interactions. Pac Symp Biocomput, 116–127

Kelly, MA, Chellgren, BW, Rucker, AL, et al. 2001 Host-guest study of left-handed polyproline ii helix formation. Biochemistry 40: 14376–14383

Kobayashi, K, Sinasac, DS, Iijima, M, et al. 1999 The gene mutated in adult-onset type ii citrullinaemia encodes a putative mitochondrial carrier protein. Nat Genet 22: 159–163

Kumar, S, Stecher, G, Li, M, Knyaz, C and Tamura, K 2018 Mega x: Molecular evolutionary genetics analysis across computing platforms. Mol Biol Evol 35: 1547–1549

Levy, ED 2010 A simple definition of structural regions in proteins and its use in analyzing interface evolution. J Mol Biol 403: 660–670

Liang, K, Li, N, Wang, X, et al. 2018 Cryo-em structure of human mitochondrial trifunctional protein. Proc Natl Acad Sci U S A 115: 7039–7044

Madeira, F, Park, YM, Lee, J, et al. 2019 The embl-ebi search and sequence analysis tools apis in 2019. Nucleic Acids Res 47: W636–W641

Mommsen, TP and Walsh, PJ 1989 Evolution of urea synthesis in vertebrates: The piscine connection. Science 243: 72–75.

Morizono, H, Caldovic, L, Shi, D and Tuchman, M 2004 Mammalian N-acetylglutamate synthase. Mol Genet Metab 81 Suppl 1: S4–11

Pedretti, A, Villa, L and Vistoli, G 2002 Vega: A versatile program to convert, handle and visualize molecular structure on windows-based pcs. J Mol Graph Model 21: 47–49

Pekkala, S, Martinez, AI, Barcelona, B, et al. 2009 Structural insight on the control of urea synthesis: Identification of the binding site for n-acetyl-l-glutamate, the essential allosteric activator of mitochondrial carbamoyl phosphate synthetase. Biochem J 424: 211–220

Powers-Lee, SG, Mastico, RA and Bendayan, M 1987 The interaction of rat liver carbamoyl phosphate synthetase and ornithine transcarbamoylase with inner mitochondrial membranes. J Biol Chem 262: 15683–15688

Qu, Q, Morizono, H, Shi, D, Tuchman, M and Caldovic, L 2007 A novel bifunctional n-acetylglutamate synthase-kinase from xanthomonas campestris that is closely related to mammalian N-acetylglutamate synthase. BMC biochemistry 8: 4

Raijman, L 1976 Enzyme and reactant concentrations and the regulation of urea synthesis. S Grisolia, R Baguena and F Mayor The urea cycle New York, London, Sydney, Toronto John Wiley & Sons, 243–259

Raijman, L and Jones, ME 1976 Purification, composition, and some properties of rat liver carbamyl phosphate synthetase (ammonia). Arch Biochem Biophys 175: 270–278

Rardin, MJ, Taylor, GS and Dixon, JE 2009 Distinguishing mitochondrial inner membrane orientation of dual specific phosphatase 18 and 21. Methods Enzymol 457: 275–287

Rardin, MJ, Wiley, SE, Murphy, AN, Pagliarini, DJ and Dixon, JE 2008 Dual specificity phosphatases 18 and 21 target to opposing sides of the mitochondrial inner membrane. J Biol Chem 283: 15440–15450

Robinson, JB, Jr., Inman, L, Sumegi, B and Srere, PA 1987 Further characterization of the krebs tricarboxylic acid cycle metabolon. J Biol Chem 262: 1786–1790

Robinson, JB, Jr. and Srere, PA 1985 Organization of krebs tricarboxylic acid cycle enzymes in mitochondria. J Biol Chem 260: 10800–10805

Rubin, GM, Yandell, MD, Wortman, JR, et al. 2000 Comparative genomics of the eukaryotes. Science 287: 2204–2215

Salka, K, Bhuvanendran, S, Wilson, K, et al. 2017 Superresolution imaging identifies that conventional trafficking pathways are not essential for endoplasmic reticulum to outer mitochondrial membrane protein transport. Sci Rep 7: 16

Schmitt, DL and An, S 2017 Spatial organization of metabolic enzyme complexes in cells. Biochemistry 56: 3184–3196

Shevchenko, A, Tomas, H, Havlis, J, Olsen, JV and Mann, M 2006 In-gel digestion for mass spectrometric characterization of proteins and proteomes. Nature protocols 1: 2856–2860

Shi, D, Morizono, H, Ha, Y, Aoyagi, M, Tuchman, M and Allewell, NM 1998 1.85-a resolution crystal structure of human ornithine transcarbamoylase complexed with N-phosphonacetyl-L-ornithine. Catalytic mechanism and correlation with inherited deficiency. J Biol Chem 273: 34247–34254

Shigesada, K and Tatibana, M 1978 N-acetylglutamate synthetase from rat-liver mitochondria. Partial purification and catalytic properties. Eur J Biochem 84: 285–291.

Shrake, A and Rupley, JA 1973 Environment and exposure to solvent of protein atoms. Lysozyme and insulin. J Mol Biol 79: 351–371

Sonoda, T and Tatibana, M 1983 Purification of N-acetyl-L-glutamate synthetase from rat liver mitochondria and substrate and activator specificity of the enzyme. J Biol Chem 258: 9839–9844.

Stankiewicz, AR, Lachapelle, G, Foo, CP, Radicioni, SM and Mosser, DD 2005 Hsp70 inhibits heat-induced apoptosis upstream of mitochondria by preventing bax translocation. J Biol Chem 280: 38729–38739

Subramanian, K, Jochem, A, Le Vasseur, M, et al. 2019 Coenzyme q biosynthetic proteins assemble in a substrate-dependent manner into domains at ER-mitochondria contacts. J Cell Biol 218: 1353–1369

Sumegi, B and Srere, PA 1984 Binding of the enzymes of fatty acid beta-oxidation and some related enzymes to pig heart inner mitochondrial membrane. J Biol Chem 259: 8748–8752

Sweetlove, LJ and Fernie, AR 2018 The role of dynamic enzyme assemblies and substrate channelling in metabolic regulation. Nat Commun 9: 2136

Tuchman, M 1989 Persistent acitrullinemia after liver transplantation for carbamylphosphate synthetase deficiency. N Engl J Med 320: 1498–1499

Velot, C, Mixon, MB, Teige, M and Srere, PA 1997 Model of a quinary structure between krebs tca cycle enzymes: A model for the metabolon. Biochemistry 36: 14271–14276

Wang, D, Eraslan, B, Wieland, T, et al. 2019a A deep proteome and transcriptome abundance atlas of 29 healthy human tissues. Mol Syst Biol 15: e8503

Wang, Y, Palmfeldt, J, Gregersen, N, et al. 2019b Mitochondrial fatty acid oxidation and the electron transport chain comprise a multifunctional mitochondrial protein complex. J Biol Chem 294: 12380–12391

Waterlow, JC 1999 The mysteries of nitrogen balance. Nutrition Research Reviews 12: 25–54

Wu, F and Minteer, S 2015 Krebs cycle metabolon: Structural evidence of substrate channeling revealed by cross-linking and mass spectrometry. Angew Chem Int Ed Engl 54: 1851–1854

Xia, C, Fu, Z, Battaile, KP and Kim, JP 2019 Crystal structure of human mitochondrial trifunctional protein, a fatty acid beta-oxidation metabolon. Proc Natl Acad Sci U S A 116: 6069–6074

Yokota, S and Mori, M 1986 Immunoelectron microscopical localization of ornithine transcarbamylase in hepatic parenchymal cells of the rat. Histochem J 18: 451–457

