## Supplemental Tables 1-4 and Figure S1 for "Mitochondrial Enzymes of the Urea Cycle Cluster at the Inner Mitochondrial Membrane"

**Table S1.** Missense sequence variants found in patients with CPS1 deficiency. Nucleotide numbers correspond to GenBank entry NM_001875. PubMed and Google Scholar were searched for reports of *CPS1* sequence variants on January 25, 2020. Table has been adapted from Yan B. et al. (2019) Frontiers in Genetics, 10: 718.

| No | Exon | Codon | Nucleotide mutation | Protein | Reference |
|---|---|---|---|---|---|
| 1 | 2 | 43 | c.128C>T | p.A43V | Häberle et al., 2011 |
| 2 | 2 | 58 | c.173G>A | p.G58D | Häberle et al., 2011 |
| 3 | 2 | 58 | c.173G>T | p.G58V | Chen et al., 2018 |
| 4 | 2 | 65 | c.194C>T | p.S65F | Häberle et al., 2011 |
| 5 | 2 | 71 | c.212T>G | p.V71G | Häberle et al., 2011 |
| 6 | 2 | 79 | c.236G>A | p.G79E | Kurokawa et al.,2007 |
| 7 | 3 | 87 | c.259C>T | p.P87S | Häberle et al., 2011 |
| 8 | 3 | 89 | c.265T>G | p.Y89D | Häberle et al., 2011 |
| 9 | 3 | 99 | c.296C>T | p.P99L | Kretz et al., 2012 |
| 10 | 3 | 123 | c.368C>T | p.S123F | Summar et al., 1998 |
| 11 | 3 | 123 | c.368C>A | p.S123T | Ali et al., 2016 |
| 12 | 4 | 108 | c.323G>A | p.G108E | Yang et al., 2017 |
| 13 | 4 | 149 | c.446T>C | p.L149S | Chen et al., 2018 |
| 14 | 5 | 160 | c.478G>A | p.A160T | Fan et al, 2019 |
| 15 | 5 | 165 | c.494A>G | p.D165G | Häberle et al., 2011 |
| 16 | 5 | 174 | c.520C>T | p.R174W | Ali et al., 2016 |
| 17 | 7 | 212 | c.634T>A | p.Y212N | Kurokawa et al., 2007 |
| 18 | 7 | 224 | c.671A>T | p.D224V | Häberle et al., 2011 |
| 19 | 7 | 233 | c.697C>T | p.R233C | Häberle et al., 2011 |
| 20 | 8 | 243 | c.728A>C | p.H243P | Häberle et al., 2011 |
| 21 | 8 | 258 | c.773G>A | p.G258E | Häberle et al., 2011 |
| 22 | 8 | 263 | c.788G>A | p.G263E | Häberle et al., 2011 |
| 23 | 8 | 265 | c.794C>T | p.P265L | Kretz et al.,2012 |
| 24 | 8 | 266 | c.796G>A | p.G266R | Chen et al., 2018 |
| 25 | 8 | 280 | c.840G>C | p.K280N | Kurokawa et al.,2007 |
| 26 | 9 | 301 | c.902G>A | p.G301E | Eeds et al.,2006 |
| 27 | 9 | 304 | c.911C>T | p.A304V | Häberle et al., 2011 |
| 28 | 10 | 317 | c.950G>A | p.G317E | Häberle et al., 2011 |
| 29 | 10 | 337 | c.1010A>G | p.H337R | Aoshima et al., 2001 |
| 30 | 10 | 341 | c.1022 T>C | p.L341S | Funghini et al.,2012 |
| 31 | 10 | 355 | c.1063A>G | p.N355D | Eeds et al., 2006 |
| 32 | 10 | 358 | c.1072G>C | p.D358H | Häberle et al., 2011 |
| 33 | 11 | 382 | c.1145C>T | p.P382L | Häberle et al., 2011 |
| 34 | 12 | 389 | c.1166A>G | p.Y389C | Eeds et al., 2006 |
| 35 | 12 | 390 | c.1169T>G | p.L390R | Eeds et al., 2006 |
| 36 | 12 | 401 | c.1201G>C | p.G401R | Häberle et al., 2011 |
| 37 | 13 | 431 | c.1291G>A | p.G431R | Häberle et al., 2011 |
| 38 | 13 | 432 | c.1295G>T | p.G432V | Häberle et al., 2011 |
| 39 | 13 | 438 | c.1312G>A | p.A438T | Häberle et al., 2011 |
| 40 | 13 | 438 | c.1312G>C | p.A438P | Kurokawa et al., 2007 |

| | | | | | |
|---|---|---|---|---|---|
| 41 | 13 | 450 | c.1348A>G | p.K450E | Häberle et al., 2011 |
| 42 | 14 | 457 | c.1370T>G | p.V457G | Funghini et al., 2003 |
| 43 | 14 | 471 | c.1412C>A | p.T471N | Pekkala et al., 2010 |
| 44 | 14 | 498 | c.1492G>C | p.A498P | Häberle et al., 2011 |
| 45 | 15 | 531 | c.1592T>A | p.V531E | Häberle et al., 2011 |
| 46 | 15 | 531 | c.1592T>G | p.V531G | Häberle et al., 2011 |
| 47 | 15 | 544 | c.1631C>T | p.T544M | Finckh et al.,1998; Häberle et al., 2011 |
| 48 | 16 | 587 | c.1759C>T | p.R587C | Häberle et al., 2011 |
| 49 | 16 | 587 | c.1760G>A | p.R587H | Kurokawa et al., 2007; Wang et al., 2011; Häberle et al., 2011 |
| 50 | 16 | 587 | c.1760G>T | p.R587L | Häberle et al., 2011 |
| 51 | 16 | 589 | c.1765G>A | p.A589T | Eeds et al., 2006 |
| 52 | 16 | 593 | c.1777G>C | p.G593R | Kurokawa et al., 2007 |
| 53 | 16 | 597 | c.1790C>T | p.S597L | Häberle et al., 2011 |
| 54 | | 600 | c.1799G>A | p.C600Y | Zhang et al.,2018 |
| 55 | 17 | 622 | c.1864G>A | p.V622M | Häberle et al., 2011 |
| 56 | 17 | 628 | c.1883G>A | p.G628D | Häberle et al., 2011 |
| 57 | 17 | 632 | c.1895T>G | p.I632R | Häberle et al., 2011 |
| 58 | 17 | 638 | c.1913G>C | p.R638P | Häberle et al., 2011 |
| 59 | 17 | 640 | c.1918G>T | p.A640S | Eeds et al., 2006 |
| 60 | 17 | 648 | c.1943G>A | p.C648Y | Häberle et al., 2011 |
| 61 | 17 | 651 | c.1951G>A | p.E651K | Kurokawa et al., 2007 |
| 62 | 17 | 653 | c.1958T>G | p.V653G | Fan et al., 2019 |
| 63 | 17 | 654 | c.1961A>T | p.D654V | Häberle et al., 2011 |
| 64 | 17 | 661 | c.1981G>C | p.G661R | Funghini et al., 2012 |
| 65 | 17 | 661 | c.1981G>T | p.G661C | Zhang et al. (2018a) PMID 30512161 |
| 66 | 18 | 674 | c.2021A>T | p.N674I | Kurokawa et al., 2007 |
| 67 | 18 | 674 | c.2022T>A | p.N674K | Häberle et al., 2011 |
| 68 | 18 | 678 | c.2033A>C | p.Q678P | Pekkala et al., 2010 |
| 69 | 18 | 698 | c.2093A>G | p.N698S | Häberle et al., 2011 |
| 70 | 18 | 716 | c.2148T>A | p.N716K | Summar et al., 1998; Eeds et al., 2006 |
| 71 | 18 | 718 | c.2153G>A | p.R718K | Häberle et al., 2011 |
| 72 | 18 | 721 | c.2162G>A | p.R721Q | Häberle et al., 2011 |
| 73 | 18 | 724 | c.2170G>C | p.A724P | Häberle et al., 2011 |
| 74 | 18 | 726 | c.2176G>A | p.A726T | Häberle et al., 2011 |
| 75 | 19 | 767 | c.2300A>T | p.D767V | Häberle et al., 2011 |
| 76 | 19 | 774 | c.2321C>T | p.P774L | Pekkala et al. [2010] |
| 77 | 19 | 780 | c.2339G>A | p.R780H | Kurokawa et al., 2007; Häberle et al., 2011 |
| 78 | 19 | 792 | c.2376G>C | p.M792I | Häberle et al., 2011 |
| 79 | 20 | 803 | c.2407C>A | p.R803S | Häberle et al., 2011 |
| 80 | 20 | 803 | c.2407C>G | p.R803G | Ono et al.,2009; Häberle et al., 2011 |
| 81 | 20 | 803 | c.2407C>T | pR803C | Häberle et al., 2011 |
| 82 | 20 | 805 | c.2413T>C | p.F805L | Eeds et al., 2006 |
| 83 | 20 | 805 | c.2414T>C | p.F805S | Häberle et al., 2011 |
| 84 | 20 | 810 | c.2429G>A | p.Q810R | Funghini et al., 2003 & Eeds et al., 2006 |
| 85 | 20 | 814 | c.2440C>T | p.R814W | Rokicki et al., 2017; Häberle et al., 2011 |

| | | | | | |
|---|---|---|---|---|---|
| 86 | 20 | 816 | c.2446T>C | p.C816R | Häberle et al., 2011 |
| 87 | 20 | 843 | c.2528T>C | p.L843S | Häberle et al., 2003 |
| 88 | 20 | 846 | c.2537C>T | p.P846L | Zhang et al., 2018 |
| 89 | 20 | 850 | c.2548C>T | p.R850C | Kurokawa et al., 2007 |
| 90 | 20 | 850 | c.2549G>A | p.R850H | Wakutani et al., 2004; Häberle et al., 2011 |
| 91 | 21 | 871 | c.2611A>C | p.T871P | Kretz et al., 2012 |
| 92 | 21 | 875 | c.2623A>G | p.K875E | Häberle et al., 2003 |
| 93 | 22 | 911 | c.2732G>A | p.G911E | Häberle et al., 2011 |
| 94 | 22 | 911 | c.2732G>T | p.G911V | Eeds et al., 2006 |
| 95 | 22 | 913 | c.2738C>T | p.S913L | Häberle et al., 2011 |
| 96 | 22 | 914 | c.2740G>C | p.D914H | Häberle et al., 2011 |
| 97 | 22 | 914 | c.2741A>G | p.D914G | Häberle et al., 2011 |
| 98 | 22 | 918 | c.2752T>C | p.S918P | Wakutani et al., 2004 |
| 99 | 22 | 932 | c.2795G>C | p.R932T | Häberle et al., 2011 |
| 100 | 23 | 949 | c.2845G>A | p.A949T | Häberle et al., 2011 |
| 101 | 23 | 958 | c.2873T>C | p.L958P | Eeds et al., 2006 |
| 102 | 23 | 959 | c.2876A>G | p.Y959C | Häberle et al., 2011 |
| 103 | 23 | 962 | c.2885A>G | p.Y962C | Häberle et al., 2011 |
| 104 | 23 | 964 | c.2891G>A | p.G964D | Funghini et al., 2012 |
| 105 | 24 | 978 | c.2933T>A | p.V978E | Häberle et al., 2011 |
| 106 | 24 | 982 | c.2944G>A | p.G982S | Eeds et al., 2006 |
| 107 | 24 | 982 | c.2945G>A | p.G982D | Kurokawa et al., 2007; Häberle et al., 2011 |
| 108 | 24 | 982 | c.2945G>T | p.G982V | Wang et al., 2011; Häberle et al., 2011 |
| 109 | 24 | 984 | c.2950T>C | p.Y984H | Häberle et al., 2011 |
| 110 | 24 | 986 | c.2957T>C | p.I986T | Häberle et al., 2011 |
| 111 | 24 | 987 | c.2959G>T | p.G987C | Häberle et al., 2011 |
| 112 | 25 | 992 | c.2975T>C | p.F992S | Häberle et al., 2011 |
| 113 | 25 | 998 | c.2993C>T | p.S998F | Eeds et al., 2006 |
| 114 | 25 | 1016 | c.3047A>G | p.N1016S | Häberle et al., 2011 |
| 115 | 25 | 1017 | c.3050C>T | p.P1017L | Häberle et al., 2011 |
| 116 | 25 | 1022 | c.3065C>T | p.T1022I | Häberle et al., 2011 |
| 117 | 25 | 1034 | c.3101A>G | p.E1034G | Häberle et al., 2011 |
| 118 | 25 | 1045 | c.3134A>G | p.H1045R | Häberle et al., 2011 |
| 119 | 26 | 1054 | c.3161T>G | p.I1054R | Rapp et al., 2001 |
| 120 | 26 | 1059 | c.3176A>G | p.Q1059R | Häberle et al., 2011 |
| 121 | 26 | 1065 | c.3194C>A | p.A1065E | Häberle et al., 2011 |
| 122 | 26 | 1089 | c.3265C>T | p.R1089C | Khayat et al., 2009; Häberle et al., 2011 |
| 123 | 26 | 1089 | c.3266G>T | p.R1089L | Summar et al.,1998; Eeds et al., 2006 |
| 124 | 26 | 1103 | c.3308A>G | p.Q1103R | Kurokawa et al., 2007 |
| 125 | 28 | 1141 | c.3422T>G | p.V1141G | Wakutani et al., 2004; Kurokawa et al., 2007 |
| 126 | 28 | 1148 | c.3443T>A | p.M1148K | Zhang et al., 2018 |
| 127 | 28 | 1155 | c.3464C>A | p.A1155E | Häberle et al., 2011 |
| 128 | 28 | 1155 | c.3464C>T | p.A1155V | Häberle et al., 2011 |
| 129 | 29 | 1167 | c.3500C>G | p.T1167R | Funghini et al., 2012 |
| 130 | 30 | 1194 | c.3582A>C | p.E1194D | Kretz et al., 2012 |

| | | | | | |
|---|---|---|---|---|---|
| 131 | 30 | 1195 | c.3584A>C | p.H1195P | Kurokawa et al., 2007 |
| 132 | 30 | 1203 | c.3607T>C | p.S1203P | Summar et al., 1998; Eeds et al., 2006 |
| 133 | 30 | 1203 | c.3608C>T | p.S1203L | Häberle et al., 2011 |
| 134 | 30 | 1205 | c.3613G>A | p.D1205N | Eeds et al., 2006 |
| 135 | 30 | 1211 | c.3632C>G | p.P1211R | Yap et al., 2019 |
| 136 | 30 | 1215 | c.3643A>G | p.I1215V | Kurokawa et al., 2007 |
| 137 | 31 | 1228 | c.3683G>A | p.R1228Q | Häberle et al., 2011 |
| 138 | 31 | 1231 | c.3691G>C | p.A1231P | Rokicki et al., 2017 |
| 139 | 31 | 1241 | c.3723C>A | p.N1241K | Kurokawa et al., 2007 |
| 140 | 32 | 1254 | c.3760A>T | p.I1254F | Ali et al., 2016 |
| 141 | 32 | 1255 | c.3765G>C | p.E1255D | Häberle et al., 2011 |
| 142 | 32 | 1262 | c.3785G>A | p.R1262Q | Häberle et al., 2011 |
| 143 | 32 | 1262 | c.3785G>C | p.R1262P | Häberle et al., 2011 |
| 144 | 32 | 1274 | c.3820G>C | p.D1274H | Häberle et al., 2011 |
| 145 | 33 | 1317 | c.3949C>T | p.R1317W | Fan et al., 2019 |
| 146 | 33 | 1327 | c.3979T>C | p.C1327R | Häberle et al., 2011 |
| 147 | 33 | 1327 | c.3980G>A | p.C1327Y | Rokicki et al., 2017 |
| 148 | 33 | 1331 | c.3991T>C | p.S1331P | Eeds et al., 2006 |
| 149 | 33 | 1333 | c.3998G>A | p.G1333E | Häberle et al., 2011 |
| 150 | 35 | 1371 | c.4112G>T | p.R1371L | Häberle et al., 2011 |
| 151 | 35 | 1378 | c.4132G>A | p.A1378T | Eeds et al., 2006 |
| 152 | 35 | 1381 | c.4142T>C | p.L1381S | Summar et al., 1998 |
| 153 | 36 | 1391 | c.4172C>T | p.T1391M | Häberle et al., 2011 |
| 154 | 36 | 1398 | c.4192C>G | p.L1398V | Häberle et al., 2011 |
| 155 | 36 | 1411 | c.4232C>T | p.P1411L | Summar et al., 1998; Eeds et al., 2006 |
| 156 | 37 | 1439 | c.4316C>T | p.P1439L | Häberle et al., 2011 |
| 157 | 37 | 1443 | c.4327A>G | p.T1443A | Eeds et al., 2006 |
| 158 | 37 | 1453 | c.4357C>T | p.R1453W | Häberle et al., 2011; Pekkala et al., 2010 |
| 159 | 37 | 1453 | c.4358G>A | p.R1453Q | Pekkala et al. , 2010 |
| 160 | 37 | 1462 | c.4385C>G | p.P1462R | Häberle et al., 2011 |
| 161 | 38 | 1491 | c.4471T>C | p.Y1491H | Summar et al., 1998 |

**Table S2.** Missense mutations of CPS1 residues with relative accessible surface area greater than 25%.

| Apo CPS1[a] | Liganded CPS1[b] | Disease Onset | Comment | Refrence |
|---|---|---|---|---|
| p.A43V | p.A43V | | | Haberle et al., 2011 |
| p.R174W | p.R174W | | Close to interface with the L2 domain, which is important for conformational change upon NAG binding | Ali et al., 2016; de Cima et al., 2015 |
| p.R233C | p.R233C | Neonatal | | Haberle et al., 2011 |
| p.K280N | p.K280N | Neonatal | Last exon base: possible donor splice site error. Undetectable or reduced liver CPS). | Haberle et al., 2011 Kurokawa et al., 2007 |
| p.A304V | p.A304V | | The p.I986T coexists within the same allele | Haberle et al., 2011 |
| p.D358H | | | Close to interface with the L2 domain, which is important for conformational change upon NAG binding | Haberle et al., 2011 de Cima et al., 2015 |
| p.Y389C | | Neonatal | Reduced $V_{max}$ and thermal stability | Diez-Fernandez et al., 2013 Eeds et al., 2006 |
| p.G401R | p.G401R | | | Haberle et al., 2011 |
| p.A438T p.A438P | | Neonatal[c]( AtoP) | Reduced $V_{max}$ | Diez-Fernandez et al., 2013 Haberle et al., 2011 Kurokawa et al., 2007 |
| | p.K450E | | Close to interface with L3 domain and juxtaposed with the Asp1025; may disrupt electrostatic interaction. | Haberle et al., 2011 de Cima et al., 2015 |
| p.T471N | | Neonatal | 100% conserved in 233 animals. Undetectable liver CPS activity. Decreased affinity for NAG. | Pekkala et al.,2010 |
| p.A498P | p.A498P | | | Haberle et al., 2011 |
| p.G628D | | | Close to interface with L2 domain, which is important for conformational change upon NAG binding | Haberle et al., 2011 de Cima et al., 2015 |
| p.A640S | | Neonatal | | Eeds et al., 2006 |
| p.R721Q | p.R721Q | | Located in the T-loop, may affect enzymatic activity. | Haberle et al., 2011 de Cima et al., 2015 |
| p.A724P | | | Middle of L1a10 helix – may affect structure of this helix and CPS1 folding. | de Cima et al., 2015 |
| | p.R780H | Neonatal | Located in the T-loop. 6% liver CPS1 activity. | de Cima et al., 2015 Kurokawa et al., 2007 Haberle et al., 2011 |
| p.R814W | p.R814W | Late onset | Close to interface with L2 domain, which is important for conformational change upon NAG binding | Haberle et al., 2011 de Cima et al., 2015 |
| p.P846L | | | | Zhang et al., 2018 |
| p.K875E | p.K875E | Neonatal | Undetectable liver CPS activity. The p.L843S coexists within the same allele. Low yield of recombinant protein possibly due to inability to fold. | Diez-Fernandez et al., 2014 Häberle et al., 2003 |
| p.I986T | | Neonatal | The change p.A304V coexists within the same allele. | Haberle et al., 2011 |
| p.G987C | | | Last exon base: possible donor splice site error. | Haberle et al., 2011 |
| p.H1045R | p.H1045R | Neonatal | Second allele, p.S913L | Haberle et al., 2011 |
| p.Q1059R | p.Q1059R | Late onset | Less than 2% liver CPS activity. May affect structure of L3 domain. | Haberle et al., 2011 de Cima et al., 2015 |
| p.R1089C; p.R1089L | p.R1089C; p.R1089L | | Corresponding mutations in *E. coli* CPS decrease enzyme activity. | Haberle et al., 2011 Yefimenko et al., 2005 Javid-Majid et al., 1996 |

| | | | | |
|---|---|---|---|---|
| p.V1141G | | Late onset | Second allele, p.R1262X. 4.8% liver CPS activity | Wakutani et al.,2004<br>Kurokawa et al., 2007 |
| p.R1228Q | p.R1228Q | | | Haberle et al., 2011 |
| | p.I1254F | | | Ali et al., 2016 |
| p.E1255D | p.E1255D | | The mutation p.E841Q affecting the corresponding *E. coli* CPS residue reduces $V_{max}$ approx. 30-fold. | Haberle et al., 2011<br>de Cima et al., 2015 |
| p.R1262Q<br>p.R1262P | p.R1262Q<br>p.R1262P | | May disrupt nucleotide binding | de Cima et al., 2015 |
| p.D1274H | p.D1274H | | | Haberle et al., 2011 |
| p.R1317W | | Neonatal | Located in the T'-loop. May affect activation of CPS1 by NAG. | Fan et al., 2019 |
| p.C1327R | | | Located in the T'-loop. The p.C1327A recombinant CPS1 had decreased $V_{max}$ and increased $K_a^{NAG}$. | Haberle et al., 2011<br>de Cima et al., 2015 |
| p.G1333E | | | Located in the T'-loop – may disrupt catalysis. | de Cima et al., 2015 |
| p.R1371L | p.R1371L | | Reduced $V_{max}$. Increased $K_a^{NAG}$ | Diez-Fernandez et al., 2015<br>de Cima et al., 2015 |
| p.P1411L | p.P1411L | Late onset | Second allele, p.Q478X. CPSI expression studies show 50% decrease in specific activity, with normal NAG activation kinetics and thermal stability. Modest $V_{max}$ effect | Diez-Fernandez et al., 2015<br>de Cima et al., 2015<br>Summar, 1998<br>Eeds et al., 2006 |
| p.P1439L | p.P1439L | | Reduced $V_{max}$. Increased $K_a^{NAG}$ | Diez-Fernandez et al., 2015 |
| p.T1443A<br>p.T1443M | | Neonatal | Increased $K_a^{NAG}$ | de Cima et al., 2015<br>Eeds et al., 2006 |
| p.Y1491H | | Late onset | Increased $K_a^{NAG}$ | Diez-Fernandez et al., 2015 |

[a]Relative solvent accessible surface area was calculated for the apo CPS1 structure 5dot.
[b]Relative solvent accessible surface area was calculated for the liganded CPS1 structure 5dou.

**Table S3.** Missense sequence variants found in patients with OTC deficiency. Nucleotide numbers correspond to GenBank entry NM_000531. PubMed and Google Scholar databases were searched for reports of *OTC* sequence variants on January 25, 2020.

| No. | Codon[a] | Nucleotide change | Amino acid change | % Enzyme activity /[%] $^{15}$N ammonia incorporation[b] | Disease presentation[c] | Reference |
|---|---|---|---|---|---|---|
| 1 | 1 | c.1A>G | p.Met1Val | | Female | Oppliger Leibundgut et al. (1995) |
| 2 | 1 | c.1A>T | p.Met1Leu | | Neonatal | Yamaguchi et al. (2006) |
| 3 | 1 | c.2T>C | p.Met1Thr | | Female | Yamaguchi et al. (2006) |
| 4 | 1 | c.3G>A | p.Met1Ile | | Female | Climent and Rubio (2002b) |
| 5 | 26 | c.77G>A | p.Arg26Gln | 0% | Neonatal | Grompe et al. (1989) |
| 6 | 26 | c.77G>C | p.Arg26Pro | | Neonatal | Yamaguchi et al. (2006) |
| 7 | 39 | c.115G>T | p.Gly39Cys | | Late | Calvas et al. (1998) |
| 8 | 39 | c.116G>A | p.Gly39Asp | | No Information | Shchlechkov et al. (2009) |
| 9 | 40 | c.118C>T | p.Arg40Cys | | Late | Oppliger Leibundgut et al. (1995) |
| 10 | 40 | c.119G>A | p.Arg40His | 6% | Late | Tuchman et al. (1994b) |
| 11 | 40 | c.119G>T | p.Arg40Leu | | Late | Cavicchi et al. (2014 |
| 12 | 41 | c.122A>G | p.Asp41Gly | | Female | Yamaguchi et al. (2006) |
| 13 | 43 | c.127C>T | p.Leu43Phe | | Female | Oppliger Leibundgut et al. (1997) |
| 14 | 44 | c.131C>T | p.Thr44Ile | | Female | Yoo et al. (1996) |
| 15 | 45 | c.133C>G | p.Leu45Val | | Female | Tuchman et al. (1998) |
| 16 | 45 | c.134T>C | p.Leu45Pro | | Neonatal | Grompe et al. (1989) |
| 17 | 47 | c.140A>T | p.Asn47Ile | | Neonatal | Tuchman et al. (1997) |
| 18 | 47 | c.140A>C | p.Asn47Thr | | Female | Yamaguchi et al. (2006) |
| 19 | 48 | c.143T>C | p.Phe48Ser | | Female | Genet et al. (2000) |
| 20 | 49 | c.145A>C | p.Thr49Pro | | Female | Yamaguchi et al. (2006) |
| 21 | 50 | c.148G>A | p.Gly50Arg | | Late | Tuchman et al. (1997) |
| 22 | 52 | c.154G>A | p.Glu52Lys | | Neonatal | McCullough et al. (2000) |
| 23 | 52 | c.155A>G | p.Glu52Gly | | Neonatal | Yamaguchi et al. (2006) |
| 24 | 52 | c.156A>T | p.Glu52Asp | 4% | Late | McCullough et al. (2000) |
| 25 | 53 | c.158T>C | p.Ile53Thr | | Late | Yamaguchi et al. (2006) |
| 26 | 53 | c.158T>G | p.Ile53Ser | | Neonatal | Yamaguchi et al. (2006) |
| 27 | 55 | c.163T>G | p.Tyr55Asp | 28% | Late | Nishiyori et al. (1998) |
| 28 | 56 | c.167T>C | p.Met56Thr | [54%] | Late | Tuchman et al. (1997) |
| 29 | 57 | c.170T>A | p.Leu57Gln | | Neonatal | Yamaguchi et al. (2006) |
| 30 | 59 | c.176T>G | Leu59Arg | | Late | Azevedo et al. (2006) |
| 31 | 60 | c.179C>T | p.Ser60Leu | | Female | Tuchman et al. (1997) |
| 32 | 62 | c.184G>C | p.Asp62His | | No Information | Shchlechkov et al. (2009) |
| 33 | 62 | c.185A>G | p.Asp62Gly | | Female | Caldovic et al. (2015) |
| 34 | 63 | c.188T>C | p.Leu63Pro | | Female | Oppliger Leibundgut et al. (1997) |
| 35 | 67 | c.200T>G | p.Ile67Arg | | Female | Yamaguchi et al. (2006) |
| 36 | 76 | c.227T>C | p.Leu76Ser | | Neonatal | Genet et al. (2000) |
| 37 | 77 | c.231G>T | p.Leu77Phe | [35%] | Late | McCullough et al. (2000) |
| 38 | 79 | c.236G>A | p.Gly79Glu | 0% | Neonatal | Tuchman et al. (1992) |
| 39 | 80 | c.238A>G | p.Lys80Glu | | Female | Schultz and Salo (2000) |

| | | | | | | |
|---|---|---|---|---|---|---|
| 40 | 80 | c.240G>T | p.Lys80Asn | | Late | Galloway et al. (2000) |
| 41 | 83 | c.248G>A | p.Gly83Asp | | Neonatal | Bartholomew & McClellan (1998) |
| 42 | 88 | c.264A>T | p.Lys88Asn | 3% | Late | Reish et al. (1993) |
| 43 | 90 | c.268A>G | p.Ser90Gly | (<20%) | Female | Takanashi et al. (2002) |
| 44 | 90 | c.269G>A | p.Ser90Asn | | Neonatal | McCullough et al. (2000) |
| 45 | 90 | c.270T>G | p.Ser90Arg | | Female | Tuchman et al. (1998) |
| 46 | 92 | c.274C>G | p.Arg92Gly | | Female | Yamaguchi et al. (2006) |
| 47 | 92 | c.275G>T | p.Arg92Leu | | Neonatal | Yamaguchi et al. (2006) |
| 48 | 92 | c.275G>A | p.Arg92Gln | 0% | Neonatal | Grompe et al. (1991) |
| 49 | 92 | c.275G>C | p.Arg92Pro | | Neonatal | Yamaguchi et al. (2006) |
| 50 | 93 | c.277A>G | p.Thr93Ala | | Late | Tuchman and Plante (1995) |
| 51 | 93 | c.278C>T | p.Thr93Ile | | Late?? Female | |
| 52 | 94 | c.281G>C | p.Arg94Thr | | Late | Tuchman et al. (1992) |
| 53 | 95 | c.284T>C | p.Leu95Ser | | Late | McCullough et al. (2000) |
| 54 | 98 | c.292G>A | p.Glu98Lys | 33% | Female | Bisanzi et al. (2002) |
| 55 | 100 | c.299G>A | p.Gly100Asp | | Female | Oppliger Leibundgut et al. (1997) |
| 56 | 100 | c.298G>C | Gly100Arg | | Late | Kim et al. (2006) |
| 57 | 102 | c.304G>C | pAla102Pro | | Neonatal | Storkanova et al. (2013) |
| 58 | 102 | c.305C>A | p.Ala102Glu | | Neonatal | Tuchman et al. (1997) |
| 59 | 105 | c.314G>T | p.Gly105Val | | Female | Yamaguchi et al. (2006) |
| 60 | 105 | c.314G>A | p.Gly105Glu | | Late | Cavicchi et al. (2014) |
| 61 | 106 | c.316G>A | p.Gly106Arg | | Female | McCullough et al. (2000) |
| 62 | 106 | c.317G>A | p.Gly106Glu | <20% | Female | Takanashi et al. (2002) |
| 63 | 106 | c.317G>T | p.Gly106Val | | Female | Yamaguchi et al. (2006) |
| 64 | 109 | c.327T>C | p.Cys109Arg | | Female | Caldovic et al. (2015) |
| 65 | 111 | c.332T>C | p.Leu111Pro | | No Information | Grompe et al. (1989) |
| 66 | 117 | c.350A>G | p.His117Arg | 18% | Late | Matsuura and Matsuda (1998) |
| 67 | 117 | c.350A>T | p.His117Leu | | Late | Tuchman et al. (1994a) |
| 68 | 125 | c.374C>T | p.Thr125Met | <1% | Neonatal | Gilbert-Dussardier et al. (1996) |
| 69 | 126 | c.377A>G | p.Asp126Gly | | Neonatal | Matsuura et al. (1994) |
| 70 | 131 | c.392T>C | p.Leu131Ser | | Late | Yamaguchi et al. (2006) |
| 71 | 132 | c.394T>C | p.Ser132Pro | | Late | Bisanzi et al. (2002) |
| 72 | 132 | c.395C>T | p.Ser132Phe | [78.6%] | Late | Gyato et al. (2004) |
| 73 | 135 | c.404C>A | p.Ala135Glu | | Late | Yamaguchi et al. (2006) |
| 74 | 136 | c.407A>T | p.Asp136Val | | Late | Yamaguchi et al. (2006) |
| 75 | 137 | c.409G>C | p.Ala137Pro | | Female | Azevedo et al. (2006)) |
| 76 | 137 | c.409G>A | p.Ala137Thr | | Female | Yamaguchi et al. (2006) |
| 77 | 139 | c.416T>C | p.Leu139Ser | | Female | Tuchman et al. (1997) |
| 78 | 140 | c.418G>C | p.Ala140Pro | | Neonatal | Yamaguchi et al. (2006) |
| 79 | 140 | c.419C>A | pAla140Asp | | No Information | Shchlechkov et al. (2009) |
| 80 | 141 | c.421C>G | p.Arg141Gly | | Female | Yamaguchi et al. (2006) |
| 81 | 141 | c.422G>A | p.Arg141Gln | 0% | Neonatal | Maddalena et al. (1988b) |
| 82 | 141 | c.422G>C | p.Arg141Pro | | Female | Tuchman et al. (1997) |
| 83 | 142 | c.425T>A | p.Val142Glu | | Late | Tuchman et al. (2002) |
| 84 | 148 | c.443T>C | p.Leu148Ser | | Neonatal | Yamaguchi et al. (2006) |

| | | | | | | |
|---|---|---|---|---|---|---|
| 85 | 148 | c.443T>G | p.Leu148Trp | | Female | McCullough et al. (2000) |
| 86 | 148 | c.444G>C | p.Leu148Phe | | Female | Komaki et al. (1997) |
| 87 | 148 | c.444G>T | p.Leu148Phe | 17% | Female | Matsuura and Matsuda (1998) |
| 88 | 151 | c.452T>G | p.Leu151Arg | | Female | Yamaguchi et al. (2006) |
| 89 | 152 | c.455C>T | p.Ala152Val | 3.70% | Late | Kogo et al. (1998) |
| 90 | 155 | c.463G>T | p.Ala155Ser | | Female | Tuchman et al. (2002) |
| 91 | 155 | c.463G>C | p.Ala155Pro | | Female | Yamaguchi et al. (2006) |
| 92 | 155 | c.464C>A | p.Ala155Glu | | Neonatal | Yamaguchi et al. (2006) |
| 93 | 158 | c.472C>T | p.Pro158Ser | | Late | Storkanova et al. (2013) |
| 94 | 159 | c.476T>C | p.Ile159Thr | 1.50% | Late | Garcia-Perez et al. (1995) |
| 95 | 159 | c.477T>G | p.Ile159Met | | Late | Ben-Ari et al. (2010) |
| 96 | 160 | c.479T>G | p.Ile160Ser | | Female | Climent and Rubio (2002b) |
| 97 | 160 | c.479T>A | p.Ile160Asn | | Neonatal | Yamaguchi et al. (2006) |
| 98 | 160 | c.479T>C | p.Ile160Thr | | Neonatal | Yamaguchi et al. (2006) |
| 99 | 161 | c.481A>G | p.Asn161Asp | | Neonatal | Genet et al. (2000) |
| 100 | 161 | c.482A>G | p.Asn161Ser | | Neonatal | Tuchman and Plante (1995) |
| 101 | 161 | c.483T>A | p.Asn161Lys | <10% | Female | Takanashi et al. (2002) |
| 102 | 161 | c.483T>G | p.Asn161Lys | <10% | Female | Takanashi et al. (2002) |
| 103 | 162 | c.484G>C | p.Gly162Arg | | Female | Yamaguchi et al. (2006) |
| 104 | 162 | c.484G>A | p.Gly162Arg | | Neonatal | Feldmann et al. (1992) |
| 105 | 162 | c.485G>A | p.Gly162Glu | | Neonatal | Yamaguchi et al. (2006) |
| 106 | 164 | c.490T>C | p.Ser164Pro | | Neonatal | Yamaguchi et al. (2006) |
| 107 | 165 | c.493G>T | p.Asp165Tyr | | Late | Genet et al. (2000) |
| 108 | 168 | c.503A>C | p.His168Pro | | Late | Yamaguchi et al. (2006) |
| 109 | 168 | c.503A>G | p.His168Arg | | Female | Vella et al. (1996) |
| 110 | 168 | c.504T>A | p.His168Gln | [69%] | Late | Tuchman et al. (1997) |
| 111 | 169 | c.505C>G | p.Pro169Ala | | Female | Tuchman et al. (2002) |
| 112 | 169 | c.506C>T | p.Pro169Leu | | Neonatal | Genet et al. (2000) |
| 113 | 169 | c.506C>A | p.Pro169His | | Female | Caldovic et al. (2015) |
| 114 | 171 | c.513G>T | p.Gln171His | | Neonatal | Ali et al. (2018) |
| 115 | 172 | c.514A>T | p.Ile172Phe | | Female | Climent et al. (1999) |
| 116 | 172 | c.516C>G | p.Ile172Met | | Neonatal | Matsuura et al. (1994) |
| 117 | 172 | c.515T>A | Ile172Asn | | Late | Ogino et al. (2007) |
| 118 | 174 | c.520G>C | p.Ala174Pro | | Female | Tsai et al. (1993) |
| 119 | 175 | c.524A>G | p.Asp175Gly | | Late | Genet et al. (2000) |
| 120 | 175 | c.524A>T | p.Asp175Val | | Female | Tuchman et al. (1997) |
| 121 | 176 | c.526T>C | p.Tyr176His | | Neonatal | Tuchman et al. (2002) |
| 122 | 176 | c.527A>G | p.Tyr176Cys | 19% | Late | Oppliger Leibundgut et al. (1996b) |
| 123 | 176 | c.527A>C | p.Tyr176Leu | | Female | Azevedo et al. (2006) |
| 124 | 178 | c.533C>T | p.Thr178Met | | Neonatal | Oppliger Leibundgut et al. (1995) |
| 125 | 179 | c.535C>T | p.Leu179Phe | | Late | Fantur et al. (2013) |
| 126 | 179 | c.536T>C | p.Leu179Pro | | Neonatal | Yamaguchi et al. (2006) |
| 127 | 180 | c.539-540AG>CC | p.Gln180Pro | <10% | Neonatal | Hübler et al. (2001) |
| 128 | 181 | c.542A>G | p.Glu181Gly | | Neonatal | Tuchman et al. (1998) |

| | | | | | | |
|---|---|---|---|---|---|---|
| 129 | 182 | c.545A>T | p.His182Leu | | Neonatal | Tuchman et al. (1994a) |
| 130 | 183 | c.547T>G | p.Tyr183Asp | | Female | Oppliger Leibundgut et al. (1997) |
| 131 | 183 | c.548A>G | p.Tyr183Cys | | Neonatal | Reish et al. (1993) |
| 132 | 186 | c.557T>C | p.Leu186Pro | | Female | Azevedo et al. (2006) |
| 133 | 188 | c.562G>C | p.Gly188Arg | 2% | Neonatal | Gilbert-Dussadier et al. (1996) |
| 134 | 188 | c.563G>T | p.Gly188Val | | Female | Climent et al. (1999) |
| 135 | 188 | c.563G>C | p.Gly188Ala | | No Information | Shchlechkov et al. (2009) |
| 136 | 191 | c.571C>T | p.Leu191Phe | 5.70% | Late | Climent and Rubio (2002b) |
| 137 | 191 | c.572T>G | p.Leu191Arg | | Neonatal | Yamaguchi et al. (2006) |
| 138 | 192 | c.576C>G | p.Ser192Arg | | Neonatal | Matsuura et al. (1993) |
| 139 | 193 | c.577T>C | p.Trp193Arg | | Female | Yamaguchi et al. (2006) |
| 140 | 193 | c.577T>G | p.Trp193Gly | | Female | Yamaguchi et al. (2006) |
| 141 | 193 | c.579G>C | p.Trp193Cys | | No Information | Shchlechkov et al. (2009) |
| 142 | 194 | c.581T>C | p.Ile194Thr | | Late | Caldovic et al. (2015) |
| 143 | 195 | c.583G>A | p.Gly195Arg | 0% | Neonatal | Tuchman et al. (1994b) |
| 144 | 196 | c.586G>A | p.Asp196Asn | | Late | Yamaguchi et al. (2006) |
| 145 | 196 | c.586G>T | p.Asp196Tyr | | Neonatal | Tuchman et al. (1998) |
| 146 | 196 | c.586G>C | p.Asp196His | | No Information | Lin et al. (2010) |
| 147 | 196 | c.587A>T | p.Asp196Val | 7% | Neonatal | Matsuura et al. (1993) |
| 148 | 197 | c.589G>A | p.Gly197Arg | | Female | Climent et al. (1999) |
| 149 | 197 | c.590G>A | p.Gly197Glu | | Female | Tuchman et al. (1998) |
| 150 | 197 | c.589G>T | p.Gly197Trp | | No Information | Shchlechkov et al. (2009) |
| 151 | 198 | c.593A>T | p.Asn198Ile | | Neonatal | Yamaguchi et al. (2006) |
| 152 | 198 | c.594C>A | p.Asn198Lys | | Female | Popowska et al. (1999) |
| 153 | 199 | c.595A>C | p.Asn199His | | Female | Ali et al. (2018) |
| 154 | 199 | c.595A>G | p.Asn199Asp | | Female | Yamaguchi et al. (2006) |
| 155 | 199 | c.596A>G | p.Asn199Ser | | Neonatal | Tuchman et al. (2002) |
| 156 | 201 | c.591C>A | p.Leu201Met d | | | Shao et al. (2017) |
| 157 | 201 | c.602T>C | p.Leu201Pro | | Neonatal | Shimadzu et al. (1998) |
| 158 | 202 | c.604C>T | p.His202Tyr | [49%] | Late | Tuchman et al. (1997) |
| 159 | 202 | c.605A>C | p.His202Pro | | Female | Staudt et al. (1998) |
| 160 | 203 | c.608C>G | p.Ser203Cys | | Female | Tuchman et al. (1994a) |
| 161 | 205 | c.613A>G | p.Met205Val | | Neonatal | Genet et al. (2000) |
| 162 | 205 | c.614T>C | p.Met205Thr | | Neonatal | Kim et al. (2006) |
| 163 | 206 | c.617T>G | p.Met206Arg | | Neonatal | Tuchman et al. (1997) |
| 164 | 206 | c.618G>C | p.Met206Ile | | Female | Climent and Rubio (2002b) |
| 165 | 207 | c.620G>A | p.Ser207Asn | | Neonatal | Yamaguchi et al. (2006) |
| 166 | 207 | c.621C>A | p.Ser207Arg | | Neonatal | Shimadzu et al. (1998) |
| 167 | 208 | c.622G>A | p.Ala208Thr | 4% | Late | van Diggelen et al. (1996) |
| 168 | 209 | c.626C>A | p.Ala209Glu | | | Bailly et al. (2015) |
| 169 | 209 | c.626C>T | p.Ala209Val | 1% [1.4%] | Neonatal | Garcia-Perez et al. (1995) |
| 170 | 210 | c.628A>C | p.Lys210Glu | | Female | Storkanova et al. (2013) |
| 171 | 210 | c.630A>C | pLys210Asn | | Female | Azevedo et al. (2006) |
| 172 | 210 | c.628A>C | p.Lys210Gln | 0% | Female | Valik et al. (2004) |
| 173 | 213 | c.637T>A | p.Met213Lys | | Female | Oppliger Leibundgut et al. (1997) |

| | | | | | | |
|---|---|---|---|---|---|---|
| 174 | 213 | c.637T>C | p.Met213Thr | | Female | Caldovic et al. (2015) |
| 175 | 213 | c.637T>G | p.Met213Arg | | No Information | Caldovic et al. (2015) |
| 176 | 214 | c.640C>T | p.His214Tyr | | Neonatal | Yoo et al. (1996) |
| 177 | 215 | c.643C>T | p.Leu215Phe | 17% | Female | Ueta et al. (2001) |
| 178 | 216 | c.646C>G | p.Gln216Glu | | Neonatal | Grompe et al. (1989) |
| 179 | 217 | c.650C>A | p.Ala217Glu | | Female | Yamaguchi et al. (2006) |
| 180 | 218 | c.653C>T | p.Ala218Val | | No Information | Shchlechkov et al. (2009) |
| 181 | 220 | c.658C>G | p.Pro220Ala | 35% | Late | Oppliger Leibundgut et al. (1996b) |
| 182 | 220 | c.658C>T | p.Pro220Ser | | | Yu et al. (2019) |
| 183 | 220 | c.659C>T | p.Pro220Leu | | Neonatal | Yamaguchi et al. (2006) |
| 184 | 221 | c.663G>C | p.Lys221Asn | | Neonatal | Yamaguchi et al. (2006) |
| 185 | 225 | c.673C>A | p.Pro225Thr | [42%] | Late | Tuchman et al. (1994b) |
| 186 | 225 | c.674C>G | p.Pro225Arg | 0% | Neonatal | Garcia-Perez et al. (1997) |
| 187 | 225 | c.674C>T | p.Pro225Leu | 0% [0.45%] | Neonatal | Hentzen et al. (1991) |
| 188 | 233 | c.698C>T | p.Ala233Val | | Neonatal | Yamaguchi et al. (2006) |
| 189 | 239 | c.716A>T | p.Glu239Val | | Neonatal | Yamaguchi et al. (2006) |
| 190 | 239 | c.716A>G | p.Glu239Gly | | Late | Yamaguchi et al. (2006) |
| 191 | 239 | c.717G>C | p.Glu239Asp | | Female | Yamaguchi et al. (2006) |
| 192 | 242 | c.725C>T | p.Thr242Ile | | Late | Tuchman et al. (1997) |
| 193 | 244 | c.731T>A | p.Leu244Gln | 8% | Late | Calvas et al. (1998) |
| 194 | 247 | c.740C>A | p.Thr247Lys | 0% | Neonatal | Tuchman and Plante (1995) |
| 195 | 249 | c.746A>G | p.Asp249Gly | | Neonatal | Kim et al. (2006) |
| 196 | 250 | c.749C>T | p.Pro250Leu | | Late | Caldovic et al. (2015) |
| 197 | 253 | c.757G>A | p.Ala253Thr | | Neonatal | Yamaguchi et al. (2006) |
| 198 | 253 | c.757G>C | p.Ala253Pro | | Neonatal | Yamaguchi et al. (2006) |
| 199 | 255 | c.764A>C | p.His255Pro | | Female | Tuchman et al. (1998) |
| 200 | 260 | c.779T>C | p.Leu260Ser | | Female | Yamaguchi et al. (2006) |
| 201 | 261 | c.782T>C | p. Ile261Thr | | Neonatal | Li et al. (2018) |
| 202 | 262 | c.785C>A | p.Thr262Lys | 26% | Late | Giorgi et al. (2000) |
| 203 | 262 | c.785C>T | p.Thr262Ile | | Late | Yamaguchi et al. (2006) |
| 204 | 263 | c.787G>A | p.Asp263Asn | | Female | Tuchman et al. (1997) |
| 205 | 263 | c.788A>G | p.Asp263Gly | | Female | Tuchman et al. (1998) |
| 206 | 264 | c.790A>G | p.Thr264Ala | 22% | Late | Matsuura et al. (1993) |
| 207 | 264 | c.791C>A | p.Thr264Asn | | No Information | Hwu et al. (2003) |
| 208 | 264 | c.791C>T | p.Thr264Ile | | Late | Shimadzu et al. (1998) |
| 209 | 265 | c.793T>C | p.Trp265Arg | | Late | Yamaguchi et al. (2006) |
| 210 | 265 | c.794G>T | p.Trp265Leu | 56% | Late | Giorgi et al. (2000) |
| 211 | 267 | c.799A>C | p.Ser267Arg | | Female | Shimadzu et al. (1998) |
| 212 | 268 | c.803T>C | p.Met268Thr | 6.70% | Late | Matsuura et al. (1993) |
| 213 | 268 | c.802A>G | p.Met268Val | | No Information | Jamroz at al. (2013) |
| 214 | 269 | c.806G>A | p.Gly269Glu | 2% | Neonatal | Zimmer et al. (1995) |
| 215 | 269 | c.805G > A | p.Gly269Arg | | Neonatal | Bijarnia-Mahay et al. (2018); Shao et al. (2017) |
| 216 | 270 | c.809A>C | p.Gln270Pro | | Female | Yamaguchi et al. (2006) |
| 217 | 277 | c.829C>T | p.Arg277Trp | 5% [59%] | Late | Finkelstein et al. (1990a) |
| 218 | 277 | c.830G>A | p.Arg277Gln | 7% | Late | Tuchman et al. (1994b) |

| | | | | | | |
|---|---|---|---|---|---|---|
| 219 | 277 | c.830G>T | p.Arg277Leu | | Late | Tuchman et al. (2002) |
| 220 | 281 | c.842T>C | p.Phe281Ser | | Neonatal | Kim et al. (2006) |
| 221 | 284 | c.850T>A | p.Tyr284Asn | | Female | Chongsrisawat V et al. (2018) |
| 222 | 287 | c.860A>C | p.Thr287Pro | | Female | Caldovic et al. (2015) |
| 223 | 289 | c.867G>C | p.Lys289Asp | | Neonatal | Caldovic et al. (2015) |
| 224 | 289 | c.867G>T | p.Lys289Asn | 0% | Neonatal | Tuchman et al. (2002) |
| 225 | 297 | c.889G>T | p.Aso297Tyr | | No Information | Shchlechkov et al. (2009) |
| 226 | 298 | c.892T>C | p.Trip298Arg | | Female | Caldovic et al. (2015) |
| 227 | 298 | c.893G>C | p.Trp298Ser | 0% | Neonatal | Ensenauer et al. (2005) |
| 228 | 301 | c.902T>C | p.Leu301Ser | | Female | Caldovic et al. (2015) |
| 229 | 301 | c.903A>T | p.Leu301Phe | 3% | Late | Climent and Rubio (2002b) |
| 230 | 302 | c.904C>T | p.His302Tyr | 0% | Neonatal | Oppliger Leibundgut et al. (1996b) |
| 231 | 302 | c.905A>G | p.His302Arg | | Neonatal | Genet et al. (2000) |
| 232 | 302 | c.905A>T | p.His302Leu | | Female | Gilbert-Dussadier et al. (1996) |
| 233 | 302 | c.906C>G | p.His302Gln | | Late | Tuchman et al. (1997) |
| 234 | 303 | c.907T>C | p.Cys303Arg | | Neonatal | Calvas et al. (1998) |
| 235 | 303 | c.907T>G | p.Cys303Gly | | Neonatal | Tuchman et al. (2002) |
| 236 | 303 | c.908G>A | p.Cys303Tyr | | Female | Tuchman et al. (1997) |
| 237 | 304 | c.912G>T | p.Leu304Phe | 6% [74%] | Late | Tuchman et al. (1992) |
| 238 | 305 | c.914C>G | p.Pro305Arg | | Neonatal | Yamaguchi et al. (2006) |
| 239 | 305 | c.914C>A | p.Pro305His | | Female | Climent and Rubio (2002b) |
| 240 | 306 | c.917G>C | p.Arg306Thr | | No Information | Meng et al. (2013) |
| 241 | 310 | c.929A>G | p.Glu310Gly | | Late | Yamaguchi et al. (2006) |
| 242 | 311 | c.931G>A | p.Val311Met | | Late | Yamaguchi et al. (2006) |
| 243 | 311 | c.932T>A | p.Val311Glu | | Neonatal | Caldovic et al. (2015) |
| 244 | 315 | c.943G>T | p.Val315Phe | | Female | Yamaguchi et al. (2006) |
| 245 | 315 | c.944T>A | p.Val315Asp | | Female | Tuchman et al. (2002) |
| 246 | 315 | c.944T>G | p.Val315Gly | | Female | Tuchman et al. (2002) |
| 247 | 316 | c.947T>C | p.Phe316Ser | | Female | Tuchman et al. (2002) |
| 248 | 318 | c.953C>T | p.Ser318Phe | | Female | Genet et al. (2000) |
| 249 | 320 | c.959G>T | p.Arg320Leu | [3.9%] | Neonatal | Grompe et al. (1991) |
| 250 | 322 | c.964C>G | p.Leu322Val | | Female | Caldovic et al. (2015) |
| 251 | 322 | c.965T>C | p.Leu322Pro | | No Information | Shchlechkov et al.((2009) |
| 252 | 323 | c.967G>A | p.Val323Met | | Late | Kim et al. (2006) |
| 253 | 326 | c.976G>A | p.Glu326Lys | | Female | Popowska et al. (1999) |
| 254 | 330 | c.988A>G | p.Arg330Gly | | Female | Tuchman et al. (1997) |
| 255 | 332 | c.994T>A | p.Trp332Arg | 0% | Neonatal | Rapp et al. (2001) |
| 256 | 332 | c.995G>C | p.Trp332Ser | | Neonatal | Wang et al. (2014) |
| 257 | 335 | c.1005G>A | p.Met335Ile | | Neonatal | Tuchman et al. (2002) |
| 258 | 336 | c.1006G>T | p.Ala336Ser | | Late | Tuchman et al. (1998) |
| 259 | 337 | c.1009G>C | p.Val337Leu | <5% | Late | Matsuda and Tanase (1997) |
| 260 | 339 | c.1015G>C | p.Val339Leu | | Neonatal | Tuchman et al. (1997) |
| 261 | 339 | c.1016T>G | p.Val339Gly | | Neonatal | Wang et al. (2014) |
| 262 | 340 | c.1018T>C | p.Ser340Pro | | Female | Oppliger Leibundgut et al. (1997) |
| 263 | 341 | c.1022T>C | p.Leu341Pro | | Female | Climent and Rubio (2002b) |

| | | | | | | |
|---|---|---|---|---|---|---|
| 264 | 343 | c.1028C>G | p.Thr343Arg | | Neonatal | Caldovic et al. (2015) |
| 265 | 343 | c.1028C>A | p.Thr343Lys | | Female | Tuchman and Plante (1995) |
| 266 | 345 | c.1033T>C | p.Tyr345His | | Late | Yamaguchi et al. (2006) |
| 267 | 345 | c.1033T>G | p.Tyr345Asp | | Female | Tuchman et al. (1992) |
| 268 | 345 | c.1034A>G | p.Tyr345Cys | | Female | Caldovic et al. (2015) |
| 269 | 347 | c.1039C>A | p.Pro347Thr | | Female | Yamaguchi et al. (2006) |
| 270 | 347 | c.1039C>T | p.Pro347Ser | | Neonatal | Caldovic et al. (2015) |
| 271 | 347 | c.1040C>T | p.Pro347Leu | | Female | Caldovic et al. (2015) |
| 272 | 349 | c.1046T>C | p.Leu349Pro | | Female | Caldovic et al. (2015) |
| 273 | 354 | c.1061T>G | p.Phe354Cys | 1.80% | Late | Myers and Shook (1996) |

[a]Nucleotide +1 is the A of the translation initiation codon of the NM_000531.3.
[b](%) residual activity in liver or intestine or determined by expression studies; [15N] residual nitrogen incorporation into urea.
[c]Neonatal – hyperammonemia within the first week of life, severe phenotype; Late – late onset, milder phenotype.
[d]Nucleotide change c.591C>A does not correspond to amino acid change p.L201M

**Table S4.** Missense mutations of OTC residues with relative accessible surface area greater than 25%.

| Mutation | Disease Onset | Comment | Reference |
|---|---|---|---|
| p.G39C | Late | | Calvas et al. [1998] |
| p.G39D | | | Shchlechkov et al. [2009] |
| p.R40C | Late | 6% residual liver activity. Impaired import of pre-OTC into mitochondria. | Oppliger Leibundgut et al. [1995] Tuchman et al. [1994a] Mavinakere M et al. [2001] Cavicchi et al. [2014] |
| p.R40H | | | |
| p.R40L | | | |
| p.D41G | Female | | Yamaguchi et al. [2006] |
| p.N47I | Neonatal | | Tuchman et al. [1997] Yamaguchi et al. [2006] |
| p.N47T | Female | | |
| p.T49P | Female | Replacement with proline can affect protein folding and stability. | Yamaguchi et al. [2006] Shi et al. (1998) |
| p.G50R | Late | | Tuchman et al. [1997] |
| p.E52K | Neonatal | The p.E52D has 4% residual liver activity. | McCullough et al. [2000] |
| p.E52G | Neonatal | | Yamaguchi et al. [2006] |
| p.E52D | Late | | McCullough et al. [2000] |
| p.I67R | Female | | Yamaguchi et al. [2006] |
| p.L76S | Neonatal | | Genet et al. [2000] |
| p.G79E | Neonatal | 0% residual liver activity. May affect assembly of the OTC trimer. | Tuchman et al. [1992] Tuchman et al. [1995] |
| p.K80E | Female | | Schultz and Salo [2000] Galloway et al. [2000] |
| p.K80N | Late | | |
| p.A102P | Neonatal | The A102P replacement can disrupt structure of the H2 a-helix. | Storkanova et al. [2013] Tuchman et al. (1997) Shi et al. (1998) |
| p.A102E | Neonatal | | |
| p.G105V | Female | | Yamaguchi et al. [2006] |
| p.G105E | Late | | Cavicchi et al. [2014] |
| p.T125M | Neonatal | <1% residual liver activity. | Gilbert-Dussardier et al. [1996] |
| p.D136V | Late | | Yamaguchi et al. [2006] |
| p.A152V | Late | 3.70% residual liver activity. | Kogo et al. [1998] |
| p.D165Y | Late | | Genet et al. [2000] |
| p.E181G | Neonatal | | Tuchman et al. [1998] |
| p.H182L | Neonatal | May have decreased stability due to loss of ionic interaction with E297. | Tuchman et al. [1994b] Tuchman et al. [1995] |
| p.Y183D | Female | | Oppliger Leibundgut et al. [1995] Reish et al. [1993] |
| p.Y183C | Neonatal | | |
| p.G188R | Neonatal | p.G188R has 2% residual liver activity. | Gilbert-Dussardier et al. [1996] Climent et al. [1999] Shchlechkov et al. [2009] |
| p.G188V | Female | | |
| p.G188A | No Info | | |
| p.K210E | Female | The p.K210Q has 0% residual liver activity. | Storkanova et al. [2013] |
| p.K210N | Female | | Azevedo et al. [2006] |
| p.K210Q | Female | | Valik et al. [2004] |
| p.K221N | Neonatal | Splice site error because c.663G>A (p.K221K) has 8% residual activity. | Yamaguchi et al. [2006] Shimadzu et al. [1998] |
| p.E239V | Neonatal | Splice site error because c.717G>A (p.E239E) causes OTCD | Yamaguchi et al. [2006] Shimadzu et al. [1998] |
| p.E239G | Late | | |
| p.E239D | Female | | |

| | | | |
|---|---|---|---|
| p.L244Q | Late | 8% residual liver activity. | Calvas et al. [1998] |
| p.D249G | Neonatal | | Kim et al. [2006] |
| p.H255P | Female | The H255P replacement can disrupt structure of the H8 a-helix. | Tuchman et al. (1998)<br>Shi et al. (1998) |
| p.G269E | Neonatal | G269 is part of the SMG motif that is important for substrate binding. | Zimmer et al. (1995)<br>Bijarnia-Mahay et al. (2018)<br>Shao et al. (2017) |
| p.G269R | Neonatal | | |
| p.Q270P | Female | Replacement with proline can affect protein folding and stability. | Yamaguchi et al. (2006)<br>Shi et al. (1998) |
| p.K289D | Neonatal | The p.K289D has 0% residual liver activity. May affect splicing because last bp of exon 8 is affected. | Caldovic et al. [2015]<br>Tuchman et al. [2002] |
| p.K289N | Neonatal | | |
| p.D297Y | | | Shchlechkov et al. [2009]<br>Tuchman et al. [1995] |
| p.R320L | Neonatal | Residual ureagenesis: 3.9%. | Grompe et al. [1991] |
| p.L349P | Female | Replacement with proline can affect protein folding and stability. | Caldovic et al. (2015)<br>Shi et al. (1998) |

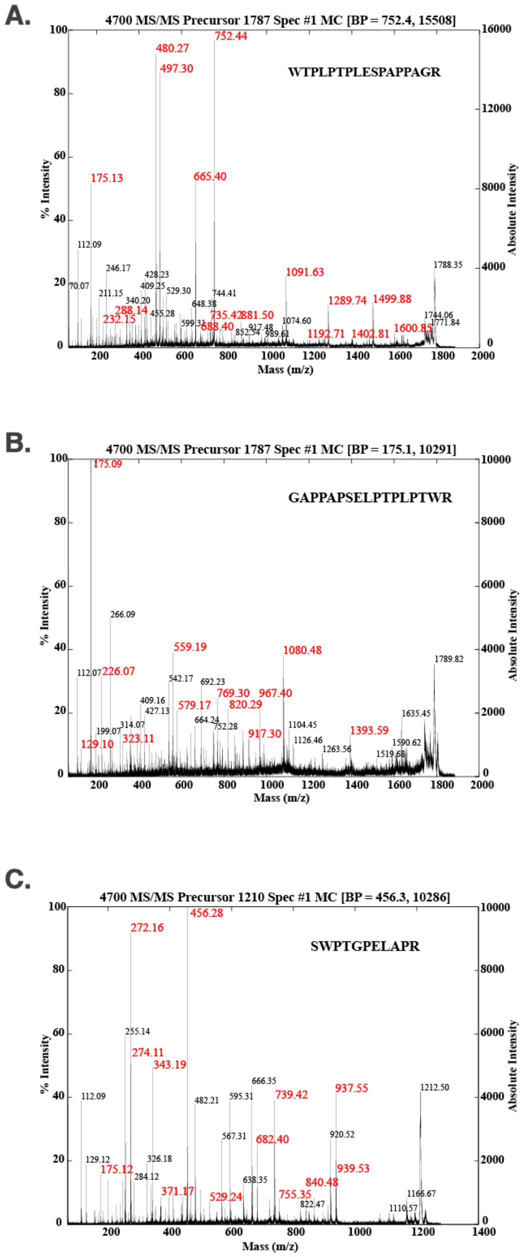

**Figure S1.** Mass spectra of purified mVS (A), rVS (B) and shVS (C).

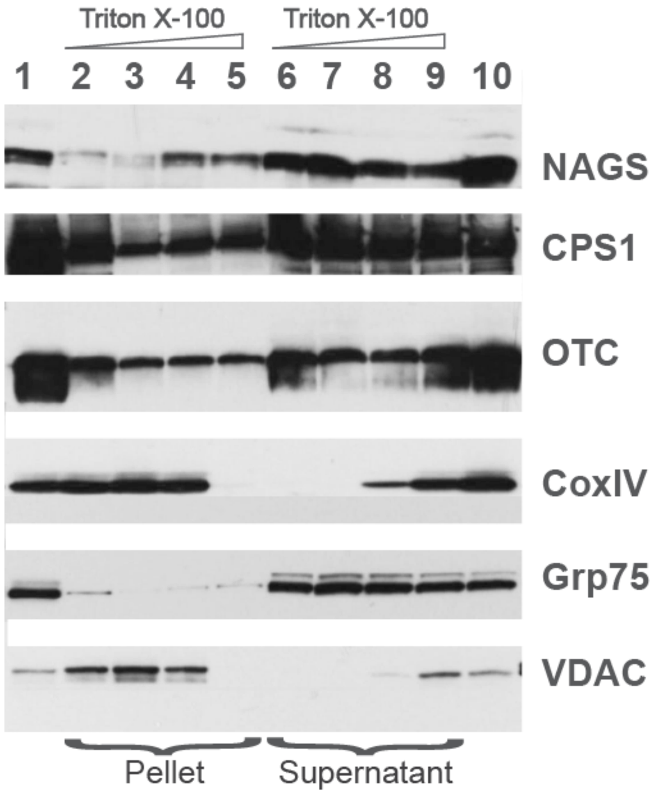

**Figure S2.** Distribution of NAGS, CPS1 and OTC in the liver mitochondria. Increasing amounts of TritonX-100 were added to sonicated mitoplasts after removal of OMM with digitonin; proteins associated with the membrane (pellet) were separated from the soluble proteins (supernatant) and probed with the anti-NAGS, anti-CPS1, anti-OTC, anti-CoxIV, anti-Grp75 and anti-VDAC antibodies. 30 μg of liver mitochondrial proteins were used as positive control for NAGS and 2 μg of mitochondrial proteins was used as positive controls for OTC, CPS1, COXIV, Grp75 and VDAC. Lane 1 – liver mitochondrial proteins, lanes 2 and 6 – 0% TritonX-100, lanes 3 and 7 – 0.1% TritonX-100, lanes 4 and 8 – 0.5% TitonX-100, lanes 5 and 9 – 1% TritonX-100, lane 10 – liver mitoplast proteins.

**References**

Ali et al. (2016) Eur. J. Pediatr. 175(3): 339–346
Ali et al. (2018) Biomed Res Int. 4320831.
Aoshima et al. (2001) Prenat. Diagn. 21(8): 634–637
Azevedo et al. (2006) Ann Hum Genet. 70(Pt 6):797-801
Bailly et al. (2015 )Neurology 85(20):e146-7
Bartholomew & McClellan (1998) Hum Mutat 12:220
Ben-Ari et al. (2010) J Hepatol. 52:292–295
Bijarnia-Mahay et al. (2018)Orphanet J Rare Dis. 13(1):174
Bisanzi et al. (2002) Mol Genet Metab 76:137–144
Caldovic et al. (2015) J Genet Genomics. 42(5):181-94
Calvas et al. (1998) Hum Mutat (Suppl 1):S81–S84
Cavicchi et al. (2014) Orphanet J Rare Dis 9, 105
Chen et al. (2018) J. Clin. Lab. Anal. 32(5): e22375
Chongsrisawat V et al. (2018) Gene 679:377-381
Climent and Rubio (2002b) Hum Mutat 19:185–186.
Climent et al. (1999) Hum Mutat 14, 352-353
de Cima et al. (2015) Sci Rep, 5: 16950
Diez-Fernandez et al. (2013) Hum Mutat. 34(8): 1149-59
Diez-Fernandez et al. (2014) Mol Genet Metab. 112(2):123-32
Diez-Fernandez et al. (2015) J Genet Genom, 42: 249-60
Eeds et al. (2006) Mol. Genet. Metab. 89(1-2): 80–86
Ensenauer et al. (2005) Mol Genet Metab 84:363–366
Fan et al. (2019) J Clin Lab Anal. E23124 [Epub ahead of print]
Fantur et al. (2013) Eur J Paediatr Neurol. 17(1):112-5
Feldmann et al. (1992) J Med Genet 29:471–475
Finckh et al. (1998) Hum. Mutat. 12(3): 206–211
Finkelstein et al. (1990) J Pediatr 117:897–902
Funghini et al. (2012) Gene 493(2): 228–234
Galloway et al. (2000) Ann Clinical Biochem 37 (Pt 5), 727-728
Garcia-Perez et al. (1995) Hum Genet 96: 549–551
Garcia-Perez et al. (1997) J Inherit Metab Dis 20:769–777.
Genet et al. (2000) J Inherit Metab Dis 23, 669-676
Gilbert-Dussadier et al. (1996) Hum Mutat 8:74–76
Gilbert-Dussardier et al. (1996) Hum Mutat 8, 74-76
Giorgi et al. (2000) Hum Mutat 15:380–381
Grompe et al. (1989) Proc Natl Acad Sci USA 86:5888–5892
Grompe et al. (1991) Am J Hum Genet 48, 212-222
Gyato et al. (2004) Ann Neurol 55:80–86.
Häberle et al. (2003) Hum. Mutat. 21(4): 444
Häberle et al. (2011) Hum Mutat. 32(6):579-89.
Hentzen et al. (1991) Hum Genet 88:153–156
Hübler et al. (2001) Z Geburtshilfe Neonatol. 205(6):236-41
Hwu et al. (2003) Hum Genet 113:365
Jamroz at al. (2013) Neurol Neurochir Pol. 47(3):283-9
Javid-Majid et al. (1996) Biochemistry 35:14362–14369
Khayat et al. (2009) Hum. Genet. 125(3): 336
Kim et al. (2006) Hum Mutat, 27, 1159
Kogo et al. (1998) J Hum Genet 43, 54-58
Komaki et al. (1997) Am J Med Genet 69: 177–181.
Kretz et al. (2012) Mol. Genet. Metab. 106 (3): 375–378

Kurokawa et al. (2007) J Hum Genet. 52(4):349-54.
Li et al. (2018) Med Sci Monit, 24: 7431–7437
Lin et al. (2010) Hum Genet. 127(4):475
Maddalena et al. (1988b) J Clin Invest 82:1353–1358.
Matsuda and Tanase (1997) Am J Med Genet 71:378–383
Matsuura and Matsuda (1998) Ryoikibetsu Shokogun Shirizu 18: 170–174.
Matsuura et al. (1993) Hum Genet 92:49–56
Matsuura et al. (1994) Hum Mutat 3:402–406.
Mavinakere M et al. (2001) J Inherit Metab Dis. 24(6):614-22
McCullough et al. (2000) Am J Med Genet 93, 313-319
Meng et al. (2013) Zhonghua Yi Xue Yi Chuan Xue Za Zhi. 30(2):195-8
Myers and Shook (1996) Am J Emerg Med 14:553–557
Nishiyori et al. (1998) Hum Mutat (Suppl 1):S131–S133
Ogino et al. (2007) Kobe J. Med. Sci. 53, 229-240
Ono et al. (2009) Brain Dev. 31(10): 779–781
Oppliger Leibundgut et al. (1995) Hum Genet 95, 191-196
Oppliger Leibundgut et al. (1996) Hum Mutat 8:333–339
Oppliger Leibundgut et al. (1997) Hum Mutat 9:409–411
Pekkala et al. (2010) Hum. Mutat. 31(7): 801-808
Popowska et al. (1999) J Inherit Metab Dis 22:92–93
Rapp et al. (2001) Eur. J. Pediatr. 160(5): 283–287
Reish et al. (1993) Biochem Med Metab Biol 50, 169-175;
Rokicki et al. (2017) Clin. Chim. Acta 47: 95–100
Schultz and Salo (2000) Arch Dis Child 82, 390-391
Shao et al. (2017) Clin Gen, 92:318–322
Shchlechkov et al. (2009) Mol Genet Metab.96(3):97-105
Shi et al. (1998) J Biol Chem 273(51): 4247–34254
Shimadzu et al. (1998) Hum Mutat (Suppl 1):S5–S7
Staudt et al. (1998) J Inherit Metab Dis 21:71–72.
Storkanova et al. (2013) Clin Genet 84, 552-559
Summar ML (1998) J. Inherit. Metab. Dis. 21 Suppl 1: 30–39
Takanashi et al. (2002) Neurology 59:210–214
Tsai et al. (1993) Hum Genet 91:321–325
Tuchman and Plante (1995) Hum Mutat 5:293–295
Tuchman et al. (1992) Pediatr Res 32, 600-604;
Tuchman et al. (1994a) Hum Mutat 4, 57-60;
Tuchman et al. (1994b) Hum Mutat 3, 318-320;
Tuchman et al. (1995) J Med Genet, 32: 680-8
Tuchman et al. (1997) J Inherit Metab Dis 20, 525-527
Tuchman et al. (1998) J Inherit Metab Dis 21(Suppl 1): 40–58.
Tuchman et al. (2002) Hum Mutat 19, 93-107
Ueta et al. (2001) Clin Chim Acta 308:187–189
Valik et al. (2004) Acta Paediatr 93, 710-711
van Diggelen et al. (1996) Clin Genet. 50:310–316.
Vella et al. (1996) J Inherit Metab Dis 20:517–524.
Wakutani et al. (2004) J Inherit Metab Dis. 27(6):787-8
Wang et al. (2011) Mol. Genet. Metab. 102(1): 103–106
Wang et al. (2014) Zhonghua Yi Xue Yi Chuan Xue Za Zhi 31, 148-151
Yamaguchi et al. (2006) Hum Mutat 27, 626-632
Yan et al. (2019) Front. Genet. 10:718
Yap et al. (2019) JIMD Rep, 48: 36-44

Yefimenko et al. (2005) J Mol Biol. 349:127–141
Yoo et al. (1996) J Inherit Metab Dis 19:31–42
Yu et al. (2019) Medicine, 98: 33
Zhang et al. (2018) J. Clin. Lab. Anal. 32(2): e222411
Zhang et al. (2018a) Zhonghua Yi Xue Yi Chuan Xue Za Zhi. 35(6):848-851
Zimmer et al. (1995) J Inherit Metab Dis 18:356–357.
